# Multi-Modal Foundation Model with Whole-Slide Attention Enables Transferrable Digital Pathology at Single-Cell Resolution

**DOI:** 10.64898/2026.07.31.741265

**Authors:** Qitian Wu, Qiyu Gong, Luezhen Yuan, Zhiyi Li, Orr Ashenberg, Fei Chen, Ramnik Xavier, Caroline Uhler

## Abstract

Paired histopathology and spatial transcriptomics data are advancing our understanding of tissue biology and disease, but modeling both modalities at single-cell resolution while mapping local and distal cell-cell interdependencies remains computationally prohibitive. Here we introduce TissueFormer, a framework for pretraining foundation models with linear rather than quadratic computational complexity, overcoming a long-standing barrier to modeling long-range dependencies at scale. Trained on over 17 million image-expression pairs from 1.2K tissue slides, TissueFormer excels at predicting spatial gene expression from histology images at cellular resolution and scales to diagnostic tasks at the cell, region, and slide levels. Additionally, by identifying both long and short-range cell-cell interdependencies, our model enables the generation of testable hypotheses about disease mechanisms and staging, as demonstrated in lung fibrosis and breast cancer.

## 1 Introduction

Recent advances in spatial biology have generated an unprecedented volume of multi-modal spatial data, particularly through the integration of histopathology images with spatially resolved transcriptomics [1–4]. These data jointly characterize tissue morphology and molecular profiles, offering transformative potential for understanding biological processes [5–8], deciphering disease mechanisms [9–12] and enhancing clinical decision-making [13–15]. Despite the rapid increase in spatial data availability, the high experimental cost—stemming from complex protocols and specialized equipment—remains a major barrier to large-scale adoption across diverse tissues, conditions, and patient cohorts [1, 2, 16]. As a result, there is increasing interest in computational approaches for cross-modality prediction, which aim to infer gene expression directly from widely available histology images [17–19]. However, the large biological and technical diversity coupled with the limited availability of paired histology and spatial transcriptomics data pose a fundamental challenge: how to extract biologically meaningful and transferable features that accommodate heterogeneity while enabling robust knowledge transfer.

Foundation models [20–23], which leverage large-scale pretraining to support diverse downstream tasks, offer a promising approach to this challenge. In computational pathology, unimodal foundation models pretrained on histopathology corpora have demonstrated strong performance in downstream predictive tasks such as slide-level and tissue-level classification or disease subtyping [24–27]. Yet, these models face important limitations in representing fine-grained, cellular-level information. More recently, bi-modal models have started to emerge that align histological features with spatial gene expression [28, 29]. However, these models operate at patch or spot resolution. As each patch aggregates hundreds of cells, this design underrepresents cell-level features critical for biological interpretation. Furthermore, while existing methods can capture observed tissue architecture through patch-level representations, they typically ignore latent intercellular dependencies, such as long-range signaling between spatially distant but functionally linked cells [30–32]. Achieving bi-modal cell-level representations and simultaneously mapping the global cell-to-cell interdependency across wholeslide images remains computationally prohibitive. Such data is inherently high-dimensional and often involves a million cells per slide. Methods that explicitly model relationships among cells must, in principle, consider interactions between every pair of cells. This quadratic growth leads to memory and computational demands that rapidly exceed the capacity of standard GPU hardware; jointly learning cell-resolved multi-modal representations and global intercellular networks at whole-slide scale is not feasible with existing architectures. Addressing this limitation requires new architectures that can integrate local and global dependencies at single-cell resolution while preserving biological interpretability [33].

In this work, we present TissueFormer, a foundation model designed for large-scale pretraining on multi-modal spatial data at single-cell resolution. To model spatial dependencies, TissueFormer introduces a whole-slide graph transformer, which captures both local dependencies between neighboring cells and global influences among arbitrary cell pairs in a tissue slide. This is achieved via a special attention mechanism that computes full all-pair interactions with linear complexity in the number of cells. Furthermore, TissueFormer incorporates a scalable contrastive loss that supports all-pair cell-level contrasts between image- and expression-based representations while avoiding quadratic computational cost.

TissueFormer is pretrained on 17 million image-expression pairs derived from over 1.2K tissue slides released by HEST-1K [34], each containing a whole-slide histology image and corresponding spatial transcriptomic profile. To evaluate the model’s generalization capacity, we establish a comprehensive benchmark comprising predictive tasks that span biological scales—from molecular profiles to cell types and diagnostic annotations. Compared to state-of-the-art histology foundation models (UNI [25], Prov-GigaPath [24], and H-optimus-1 [35]) and bi-modal foundation models (OmiCLIP [29] and GHIST [28]), TissueFormer demonstrates superior generalization to unseen tissues, species and gene markers. This enables a scalable and cost-effective framework for “virtual spatial transcriptomics”—predicting spatial gene expression at cellular resolution from routine histology images. To further assess the generalizability of the predictive capabilities of TissueFormer on tissue samples outside the HEST-1K dataset, we fine-tuned it with an independent dataset [36] and demonstrated strong performance across a wide range of tasks, including patient stratification, region of interest (ROI) disease state inference, disease-associated cell identification, pathology feature annotation, and spatial niche classification. Beyond predictive tasks, TissueFormer’s interpretable attention mechanism enables cell–cell interaction analysis directly from histology images, as demonstrated through two case studies involving tissue samples from pulmonary fibrosis [36] and breast cancer [37], respectively. In the fibrosis case, the model reveals long-range intercellular dependencies associated with early-stage disease progression. In the cancer case, its cell-level attention maps support the discovery of fine-grained disease-associated cell subtypes, providing biologically meaningful features for spatial analysis and diagnosis. Overall, TissueFormer is a powerful tool to perform virtual spatial transcriptomics at scale. Its interpretable, high-resolution maps of both local and long-range cell dependencies can provide insights into tissue organization and disease dynamics relevant for disease subtyping.

## 2 Results

### TissueFormer: A Multi-Modal Cellular-Resolution Histology Foundation Model

During pretraining, TissueFormer takes as input a haematoxylin and eosin (H&E)-stained tissue slide and its corresponding spatial transcriptomic profile (Fig. 1a). To effectively leverage the multi-modal spatial information—including histological features, gene expression, and spatial coordinates—we represent each tissue slide as a spatial adjacency graph, where each node corresponds to an individual cell (or spot) and edges reflect spatial adjacency. Each node is characterized by a cell-centered 224 × 224-pixel (with resolution scaled to be 0.5*µm*/pixel across all images) image patch extracted from the whole-slide H&E image and a cell-level gene expression vector extracted from the spatial transcriptomic profile.^1^ These image patches and gene expression vectors are processed by a histology encoder and an expression encoder that produce image-based and expression-based cell representations (i.e., numerical vectors in latent space), respectively. For expression-based representations, given that most spatial transcriptomic technologies do not provide whole-genome readouts and the set of measured genes varies across cases, we compile a unified panel consisting of all protein-coding genes measured across the dataset, spanning multiple technologies. Then the two-view representations (i.e., image- and expression-based) are aligned via a scalable multi-modal contrastive learning strategy (Fig. 1a, Methods). With inputs unified to a standard image resolution (0.5µm/pixel) and a full protein-coding gene panel, this framework is highly flexible and can be readily applied to model paired multi-modal spatial data, regardless of the measured gene set or image spatial resolution.

**Fig. 1.**
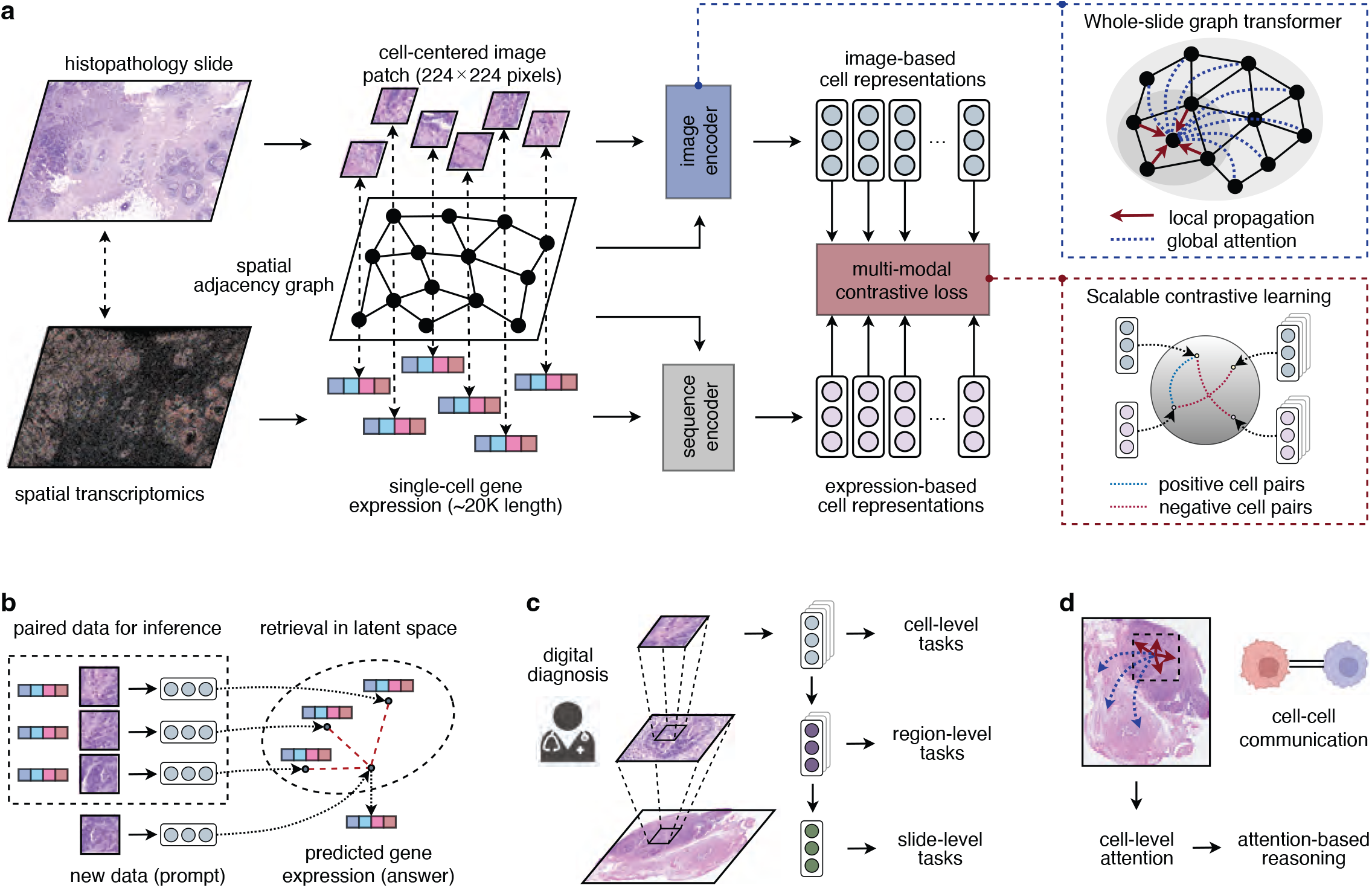
Overview of TissueFormer framework. **a**, During pretraining, TissueFormer takes an haematoxylin and eosin (H&E)-stained image (i.e., histopathology slide) paired with spatial transcriptomics as input, and represents the spatial information of a tissue slide as a cell-cell adjacency graph, where each cell is characterized by two-fold features: a cell-centered image patch extracted from the H&E image, and 2) a single-cell gene expression vector derived from spatial transcriptomics profiles. The histology and expression encoders map these multi-modal features into cell-level representations in the latent space. The histology encoder is designed as a whole-slide graph transformer which computes cell embeddings via local propagation within cell neighborhood and global attention over cells from a whole slide. The model pretraining employs contrastive learning, treating the image- and expression-based representations of a cell as positive pairs and the representations of different cells as negative pairs. **b-d**, Schematic representations of TissueFormer capabilities. For cross-modality prediction (**d**), the model leverages the paired data in the training set as reference to predict gene expression from unseen H&E images. Specifically, for any new tissue slide that only has an H&E staining, the model uses the image-based cell representations for similarity-based query (i.e., in-context learning) over the single-cell gene expression in the training set to predict cell-level gene expression of the new tissue slide. The model can be applied for informing digital diagnosis (**c**) by computing cell-, region-, and slide-level representations that can be further used for predictive tasks at different biological scales. The attention mechanism of the whole-slide graph transformer can be used to model cell-cell communication (**d**), where the cell-level attention serves as deep features uncovered in a fully data-driven manner.

Two core innovations of TissueFormer are (1) a scalable whole-slide graph transformer, and (2) a scalable contrastive learning loss, which together enable efficient spatial modeling and training at scale. The whole-slide graph transformer, implemented within the histology encoder, integrates local propagation within cell neighborhoods and global attention among cells of a whole slide to compute cell representations. Local propagation models interactions between spatially adjacent cells, while global attention captures long-range dependencies between arbitrary cells across the entire tissue slide. In contrast to existing histology foundation models (e.g., Prov-GigaPath [24]), whose attention mechanisms operate at the patch level—where each image patch aggregates features from roughly hundreds of cells—our approach performs attention directly at the cell level. This enables the construction of cell-to-cell attention maps that more accurately capture fine-grained spatial dependencies that are obscured in patch-based models (fig. S1). However, high-resolution slides can contain a million cells, making standard softmax attention [38] computationally prohibitive due to its quadratic complexity in the number of cells. To overcome this limitation, we introduce a novel attention mechanism that reduces both time and memory complexity to linear scaling by reordering matrix operations (Methods; fig. S2a,b). Importantly, by achieving linear computational complexity with respect to the number of cells (fig. S3a), this global attention mechanism preserves the ability to model dependencies between any pair of cells across the whole slide.

For contrastive learning, the image- and expression-based representations of the same cell are treated as a positive pair, while representations from different cells serve as negative pairs. Standard contrastive objectives such as the InfoNCE-based loss [39, 40] require exhaustive pairwise comparisons, resulting in quadratic computational cost. To enable efficient learning, we introduce a scalable contrastive loss that supports effective all-pair contrast at the cell level while reducing complexity to linear order (Methods; fig. S4c,d, fig. S3b). These two components—the scalable global attention and the scalable contrastive loss—together allow TissueFormer to efficiently learn informative patterns from high-resolution, multi-modal spatial data at single-cell resolution across the whole slide.

Following pretraining, TissueFormer supports a broad range of downstream applications (Fig. 1b–d). It enables “virtual spatial transcriptomics”—predicting cell-level gene expression directly from histology images using in-context learning (Fig. 1b). Moreover, it informs digital diagnosis tasks by providing rich multi-scale representations for cell-, region-, and slide-level predictions (Fig. 1c). Finally, TissueFormer ‘s interpretable attention mechanism supports *in silico* analysis of intercellular communication by modeling long-range dependencies across entire tissue slides (Fig. 1d), extending beyond the local constraints of traditional proximity-based analyses.

### TissueFormer’s predictions generalize across tissues and species

As a first evaluation, we considered cross-modality prediction, aiming to infer gene expression directly from widely available histology images (Fig.2a). We formulated this task as a high-dimensional regression problem and resolved it through an in-context learning approach which makes predictions by retrieving similar image–expression pairs from the latent space. (Methods). We demonstrated the model with a large and diverse corpus of tissue slides from HEST-1K [34], spanning 21 organs, two species (Homo sapiens and Mus musculus), four spatial transcriptomics platforms (Visium, Visium HD, ST^2^, and Xenium), and a mixture of healthy and diseased samples covering 25 cancer types (Fig. 2b). The dataset spans both sequencing-based technologies (Visium, Visium HD, ST), which produce spot-level expression profiles with large, overlapping gene panels (∼ 1.6M spots), and the imaging-based technology Xenium, which provides cell-level expression but with smaller gene panels (∼ 15M cells) (Fig. 2b). To investigate the model’s scalability, we assessed the computational efficiency of TissueFormer by measuring inference time and memory usage across Xenium slides with varying cell numbers (Fig. 2c). Even for slides with over one million cells, TissueFormer’s inference was completed in approximately one second, demonstrating its desired efficiency. Importantly, both time and memory consumption increased linearly with the number of cells per slide, confirming the model’s scalability for large-scale training and inference.

**Fig. 2.**
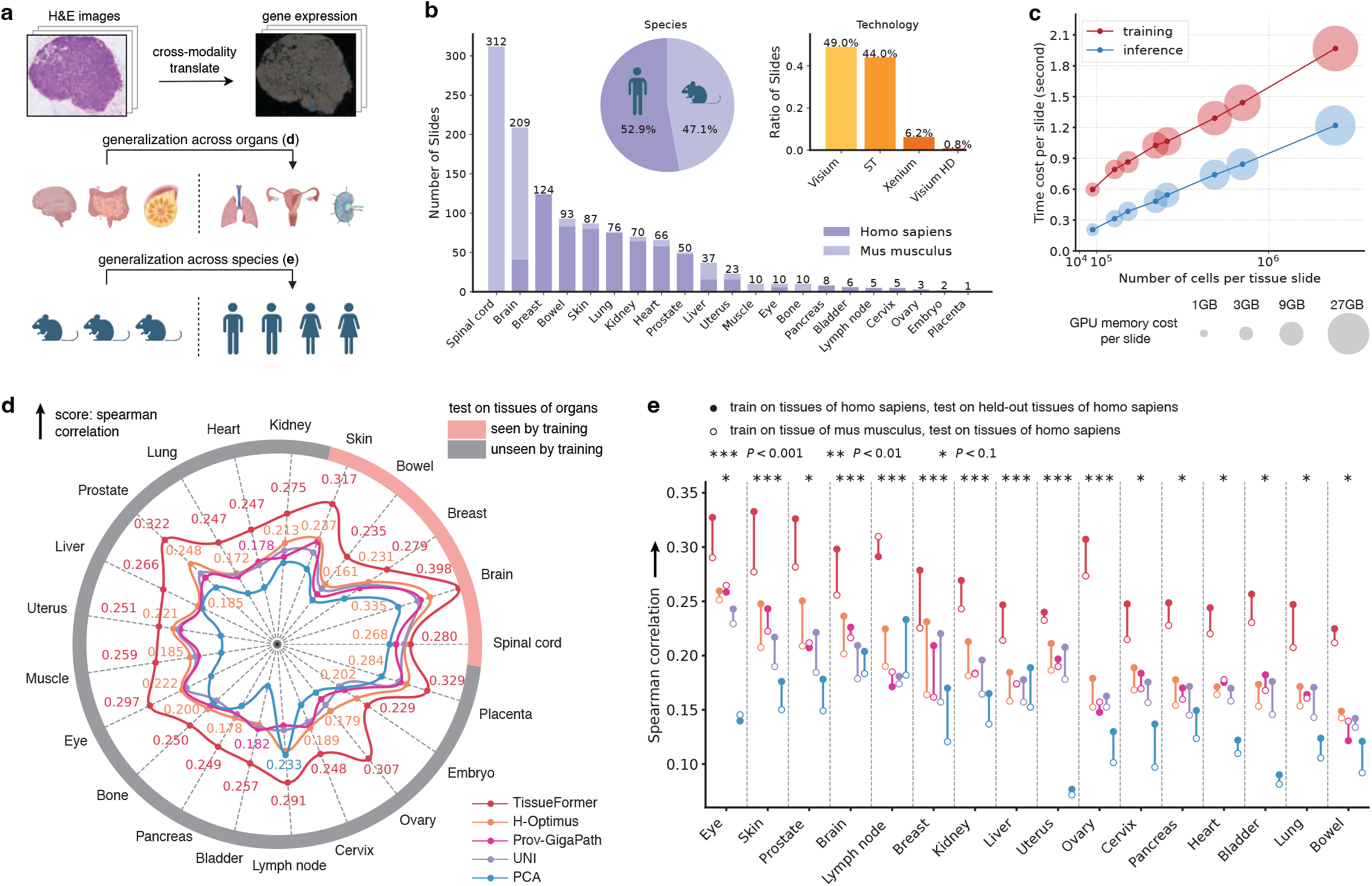
Generalization of gene expression prediction across tissues, organs and species. **a**, Illustration of the problem setting for gene expression prediction: for any test tissue slide that only has an H&E image, the model aims to predict the spot-level or cell-level gene expression of the whole slide. To evaluate the generalization capability of models, we consider two challenging evaluation protocols: 1) generalizable across organs, where tissue slides are split by organs; 2) generalization across species, where tissue slides are split by species. **b**, Statistics of the dataset from HEST-1K [34] including over 1.2K tissue slides across disparate organs, species and spatial technologies. **c**, Scalability test of TissueFormer with time and memory costs of training (time is averaged across multiple epochs) and inference on Xenium tissue slides entailing different numbers of cells ranging from 10K to 1M. **d**, Performance comparison with state-of-the-art models under the protocol of generalization across organs. Models are trained with tissue slides of the top five frequent organs in (**b**) and tested on held-out tissue slides of these organs, as well as other organs that are unseen by training. Performance is measured by spearman correlation averaged across test tissue slides: the correlation is first computed per spot between the ground-truth and predicted expression of top 4K highly variable genes and then averaged across spots of one slide. **e**, Performance comparison with state-of-the-art models under the protocol of generalization across species. The models are colored in the same way as in (**d**). We compared two specific settings: 1) models are trained with tissue slides from *homo sapiens* and tested on held-out *homo sapiens* slides; 2) models are trained with tissue slides from *mus musculus* and tested on the same *homo sapiens* slides as the first setting. The lines connecting two dots highlight the generalization gaps of different models when transferring from human to mouse tissues. The P-values indicate the significance level at which TissueFormer outperforms the best competitor in each case, based on the Wilcoxon signed-rank test.

For this initial evaluation, we focused on sequencing-based slides, whose broader gene coverage allows us to test cross-tissue generalization. We designed two evaluation protocols: one testing generalization across organs, and the other across species, by splitting training and test tissue slides by organ or species (Fig.2a). We compared TissueFormer with three state-of-the-art histology foundation models (H-optimus-1, Prov-GigaPath, UNI) and a baseline PCA model. For all models, we trained a linear regression layer to map image patch representations to gene expression. During testing, only H&E images from the test slides were used for prediction; spatial transcriptomics data were held out entirely.

To evaluate generalization across organs, we trained models on the five most frequent organs and evaluated them on both held-out slides from these organs and slides from unseen organs (Fig. 2d). We quantified performance using Spearman correlations between predicted and true expression levels of the top 1K, 2K, and 4K most variable genes, averaged across spots and slides (Fig. 2d and fig. S5). TissueFormer consistently outperformed all baselines, achieving improvements ranging from 4.2% (spinal cord) to 46.3% (bowel) in correlation scores on the five organs seen during training, and improvements ranging from 13.7% (uterus) to 71.6% (ovary) on 16 unseen organs, compared to the best competitor. Additionally, we compared our model with two recent multimodal foundation models, OmniClip [29] and GHIST [28], in this setting. The results show that TissueFormer consistently outperforms both methods across 21 different organs, as measured by Spearman correlation for the top 1K, 2K, and 4K most variable genes (fig. S6). Ablation studies further demonstrate that the global attention mechanism is the primary driver of our model’s performance gains, contributing more substantially than other components such as non-linearity and local propagation (fig. S7).

To evaluate generalization across species, we trained models on tissue slides from *Mus musculus* and tested on held-out human (*Homo sapiens*) slides, as compared to the models trained with human slides and tested on the same held-out human slides (Fig. 2e and fig. S8). As expected, all models exhibited degraded performance due to domain shift between species. Nonetheless, TissueFormer maintained robust predictive accuracy, with improvements ranging from 9.7% (eye) to 74.0% (ovary) over the best competitor. These results highlight TissueFormer ‘s strong transferability across tissues and species.

### TissueFormer accurately predicts single-cell gene expression from histology images

The ability to perform cross-modality prediction enables TissueFormer to conduct virtual spatial transcriptomics, inferring gene expression profiles directly from histology images. To evaluate the accuracy of these predictions—particularly at high spatial resolution where subcellular-level transcriptomic data are available—we introduced a gene-wise evaluation framework at single-cell resolution. We trained TissueFormer using all slides from sequencing-based technologies and a subset of Xenium imaging-based slides, and evaluated its generalizability on held-out Xenium slides and Xenium slides from an independent dataset [36] (Fig. 3). This setup enabled rigorous assessment of how well TissueFormer recapitulates spatial gene expression patterns at the cellular level, a critical factor for interpretability and downstream applications.

**Fig. 3.**
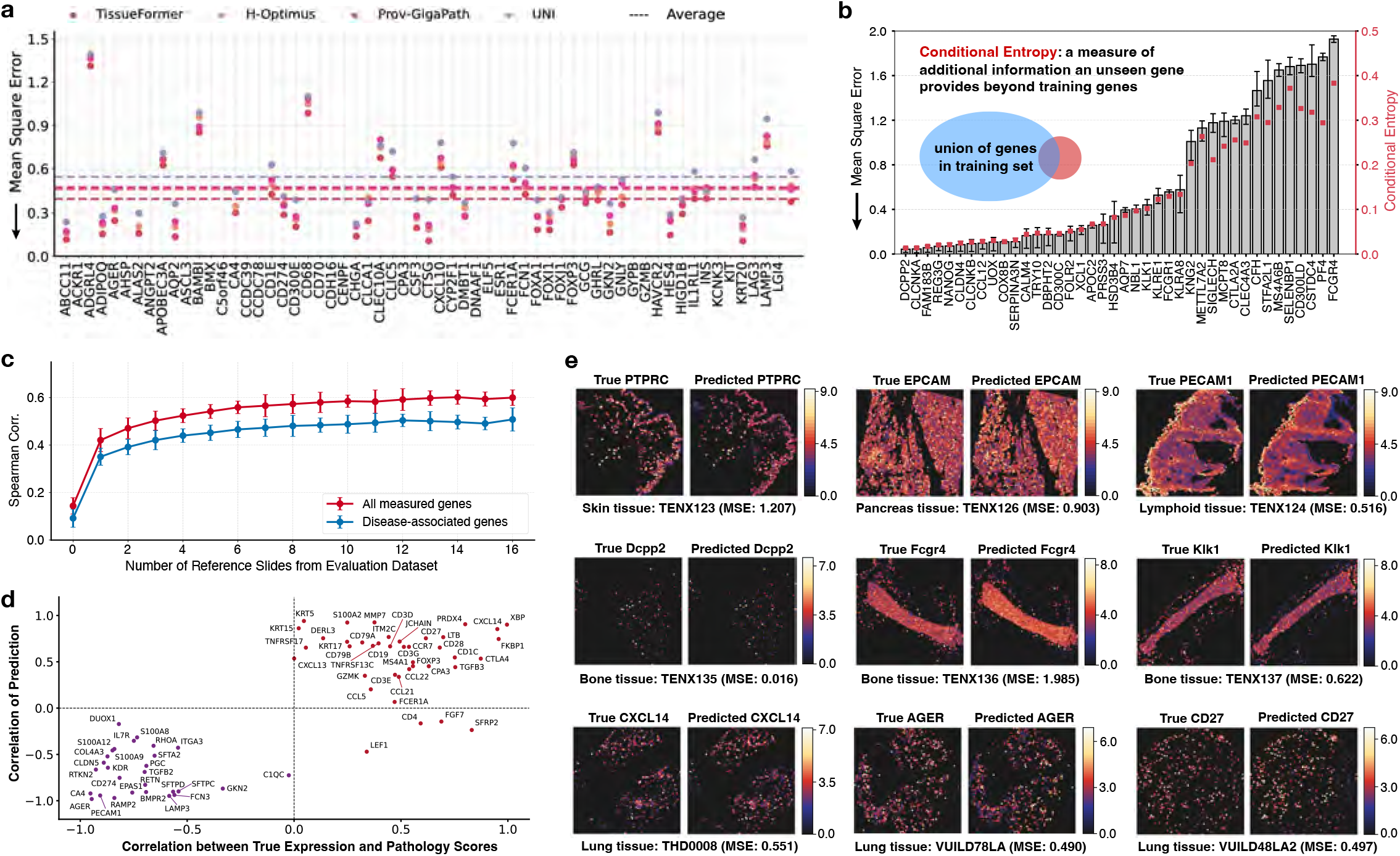
Gene-wise evaluation of gene expression prediction at single-cell resolution. **a**, Comparison of gene-wise mean square error (MSE) between ground-truth expression and expression predicted by TissueFormer and competing methods across top 60 highly-variable genes in test Xenium slides from different organs. The dotted horizontal lines show the average MSE of different models across all testing genes (TissueFormer: 0.396, H-Optimus: 0.462, Prov-GigaPath: 0.472, UNI: 0.544). **b**, Plot showing MSE between predicted and ground-truth expression of 40 new genes unseen in training. The MSE is first computed over cells per tissue slide and the error bars are computed across tissue slides in the test set. In red, we overlaid the conditional entropy level per each gene, a measure of how much additional information is provided by a new gene beyond the union of genes in the training set. **c**, Plot showing Spearman correlation between predicted and ground-truth gene expression on an independent evaluation dataset (pulmonary fibrosis [36]) when using different numbers of reference slides from the evaluation dataset. For the case with number = 0, slides from the pretraining dataset were used for reference. **d**, Plot showing the consistency of predicted gene expression with the ground-truth with respect to disease-associated markers, genes that exhibit significantly (*P <* 0.01) positive or negative correlation with pathology scores of lung tissues with pulmonary fibrosis. The Pearson correlation between ground-truth expression and pathology scores (x-axis) is compared with the Pearson correlation between predicted expression and pathology scores (y-axis). **e**, Representative images comparing the predicted and true expression levels for marker genes on tissue slides from the test set. On the first row: marker genes for common cell types; second row: marker genes from bone tissues that are unseen in training; third row: marker genes strongly associated with pulmonary fibrosis.

To help the model learn effectively across spatial resolutions and generalize better to new data, we adopted a two-step learning strategy (Methods): the model was first trained on over one thousand lower-resolution sequencing-based slides, followed by further training on several dozen high-resolution Xenium slides. We evaluated performance using gene-wise mean squared error (MSE) across the 100 most variable genes on the held-out Xenium slides (Fig. 3a, fig. S9). TissueFormer ranked first for 97 out of 100 genes and achieved an average MSE of 0.396, corresponding to a 14.5% reduction compared to the runner-up, H-optimus-1 (with an average MSE of 0.463). As a further analysis, we compared against a model trained only on sequencing-based slides (i.e., the checkpoint after the first training stage). While this version of TissueFormer underperforms the model trained on both sequencing-based and Xenium slides, it still outperforms H-Optimus and Prov-GigaPath in predicting gene expression on the held-out Xenium slides (fig. S10).

Importantly, TissueFormer’s in-context learning framework, which operates as a form of few-shot learning, enables it to generalize to the prediction of target genes by retrieving expression data from a limited number of reference slides without requiring model retraining. To evaluate this, we tested the model on three bone tissue slides, which included the expression of 40 genes randomly held-out from the training set and ranked performance by gene-wise MSE (Methods; Fig. 3b). Performance on unseen genes was negatively correlated with their conditional entropy, which measures the additional information an unseen gene provides beyond those seen in training. This trend offers insights into the model’s generalization capacity and provides a useful indicator of prediction reliability in practice.

We further tested TissueFormer on an independent dataset of pulmonary fibrosis [36], assessing its ability to generalize to new datasets without retraining. Using only three Xenium slides as reference for in-context learning, TissueFormer achieved good predictive performance on held-out test slides (with Spearman correlation above 0.5) (Fig. 3c). We then focused on gene markers that are significantly associated with pulmonary fibrosis. For each marker, we compared the Pearson correlation between predicted expression and pathology scores with that of the ground-truth expression (Fig. 3d, fig. S11). Among the 40 positively correlated and 28 negatively correlated markers, TissueFormer correctly recovered the direction of correlation for 36 and 28, respectively—demonstrating consistency with biological signal. As qualitative evidence, we visualized TissueFormer ‘s predictions alongside ground-truth expression, projected onto spatial maps of whole-slide images (Fig. 3e, fig. S12). Together, these results validate TissueFormer ‘s ability to generate spatially resolved gene expression at single-cell resolution from standard histology images, offering a scalable, cost-effective alternative to experimental spatial transcriptomics.

### TissueFormer informs digital diagnosis at cell-, region- and slide-level from histology

Histology-based diagnosis remains a cornerstone of clinical pathology, enabling experts to stratify patients by disease state, localize pathological regions, and identify cellular features indicative of disease progression [13, 41, 42]. These diagnostic tasks span multiple biological scales—from single cells to regions and entire slides—and can be naturally formulated as machine learning problems (Fig. 4a). By leveraging the pretrained TissueFormer, we aimed to address these diagnostic tasks based on just routine histology images as input.

**Fig. 4.**
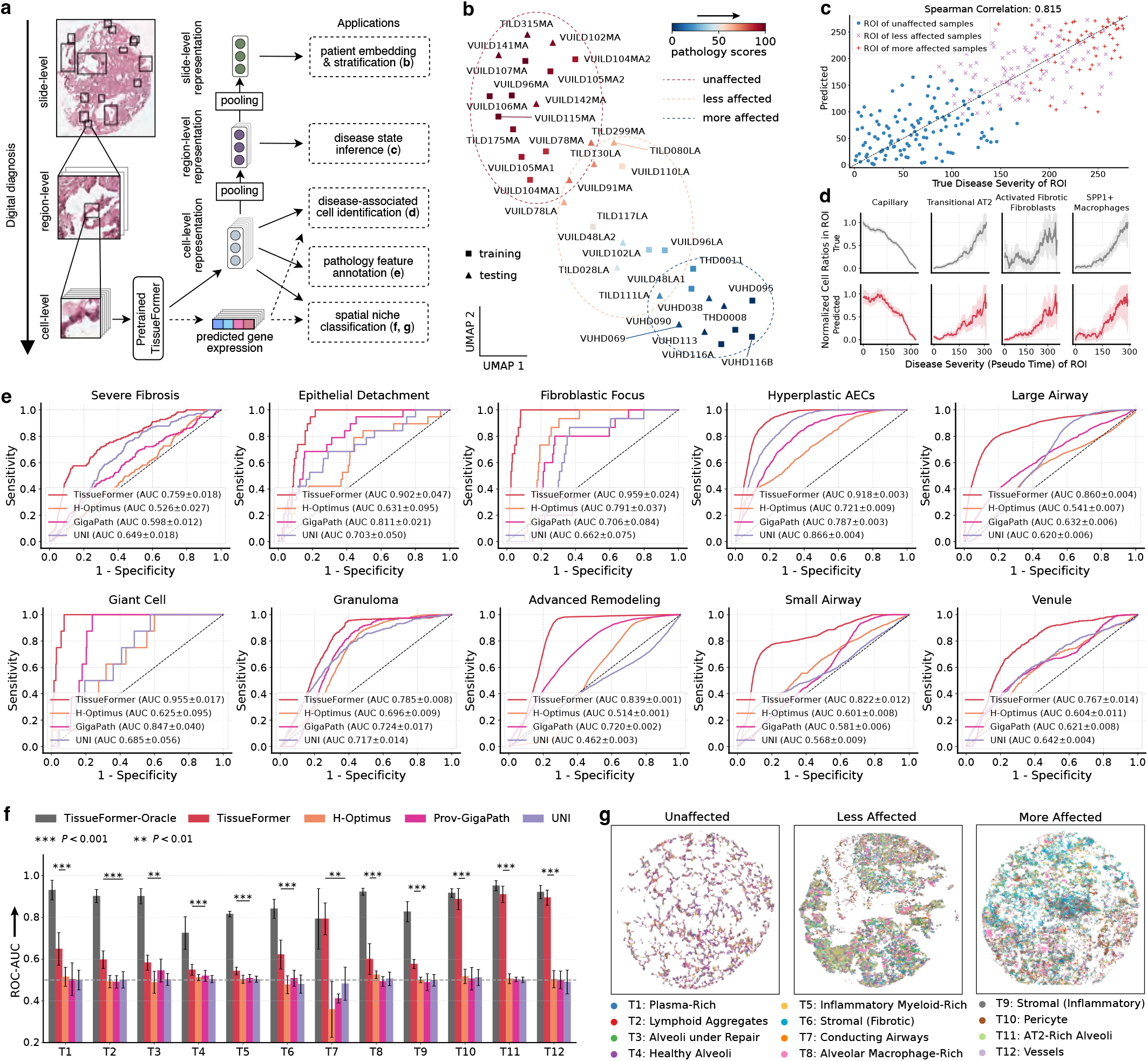
Multi-level prediction for digital pathology at different biological scales. **a**, Illustration of how the pretrained TissueFormer model is applied to multiple downstream tasks on lung tissue slides from patients with pulmonary fibrosis. The pretrained TissueFormer can handle multi-level tasks on test samples using only H&E as input. **b**, UMAP embeddings of slide-level representations that aggregate cell-level representations of a whole slide. Each point is a slide sample colored by its pathology score and the embeddings form clusters that align with the disease status (unaffected, less affected and more affected). **c**, Plot showing the correlation between the predicted and ground-truth disease severity of regions of interest (ROIs). The disease severity is a continuous label derived from the ranking of ROIs from healthy to diseased on test slides. We utilized the region-level representations by TissueFormer to infer ROIs’ disease severity. **d**, Comparison of true (top row) and predicted (bottom row) normalized cell ratios within airspace regions (ROIs) for four cell types at different stages of pulmonary fibrosis (measured by ROIs’ pseudo time from healthy, to early-stage and end-stage fibrosis) on test slides. TissueFormer uses cell-level representations from H&E images to predict cell types, and the predicted cell types are then aggregated to estimate cell type ratios within each ROI. **e**, Receiver Operating Characteristic (ROC) curves for predicting 10 annotated pathology features from different ROIs on H&E of test slides, using TissueFormer compared to competing methods. **f**, ROC-AUC scores for spatial niche classification on test slides. The cell-level representations given by TissueFormer are used to classify cells into 12 spatial niches. TissueFormer uses the predicted gene expression as input of the expression encoder to obtain predicted gene expression-based representations which are then used for prediction. TissueFormer-Oracle uses the ground-truth gene expression, acting as an oracle model providing the performance upper bound. The horizontal dotted line marks ROC-AUC of 0.5 that corresponds to uninformative random guess. The P-values indicate the significance level at which TissueFormer outperforms the best competitor in each case, based on Wilcoxon signed-rank test. **g**, Representative examples of spatial niches classified by TissueFormer on test tissue slides from unaffected, less affected and more affected pulmonary fibrosis samples.

We evaluated TissueFormer on the pulmonary fibrosis dataset comprising 36 Xenium distal lung tissue slides categorized into unaffected, less affected, and more affected groups [36]. Data were randomly split at the patient level into training and test sets. The pretrained TissueFormer was finetuned using a contrastive loss computed on the training slides. For inference, region- and slide-level representations were extracted via hierarchical pooling over the cell-level embeddings, and a shallow MLP was trained to map these representations to classification or regression targets (Methods).

We first assessed whether slide-level representations derived from H&E images could stratify patients by disease severity. UMAP projection of these representations revealed coherent clustering, where patients with similar pathology scores grouped closely (Fig. 4b), indicating that Tissue-Former encodes biologically relevant disease features. To further investigate into the heterogeneity of slides, we split each slide into four cropped sections and found the embeddings of sections from the same slide exhibit overall consistency in the UMAP space (fig. S13). Next, we tackled a region-level regression task: predicting disease severity of airspace regions, also referred to as regions of interest (ROIs). These disease severity labels, represented as continuous scores, were putatively annotated in the original study [36] based on gene expression signatures and cell-type composition. Using only H&E images, TissueFormer achieved strong agreement with the molecularly informed annotations (Fig. 4c), yielding a Spearman correlation of 0.815 on testing data. We then investigated a more challenging cell-level task: estimating the abundance of disease-associated cell types within ROIs—capillary, transitional Alveolar Type 2 (AT2), activated fibrotic fibroblasts, and SPP1^+^ macrophages—each associated with different stages of pulmonary fibrosis ^3^. We formulated this task as a binary classification problem at the cell level, and used TissueFormer ‘s cell-level embeddings to predict whether a given cell belonged to a target cell type. Aggregating predictions across cells in each ROI provided estimated cell type ratios, which closely matched the cell type ratios computed by molecularly informed annotations and recapitulated disease progression trends (Fig. 4d).

We further evaluated TissueFormer on two additional cell-level classification tasks: (1) identifying pathology annotations that represent histological features (e.g., cellular or regional morphology) associated with pulmonary fibrosis, and (2) classifying spatial niches representing certain functional subgroups. For the pathology feature annotation task, TissueFormer outperformed all baselines across ten different features, with an average ROC-AUC improvement of 40.1% over the runner-up (Fig.4e). We observed that the improvements yielded by TissueFormer were more pronounced on the pathology annotations related to cellular morphology (e.g., giant cell and fibroblastic focus) than the annotations that pertain to regional features (e.g., severe fibrosis and granuloma). This difference may stem from TissueFormer’s fine-grained pretraining at single-cell resolution that leads to better capability for capturing microscopic features. We then tested the model on the more difficult spatial niche classification task. To this end, we used the transcript-based niches defined in the original paper [36], which used only transcript data without relying on cell segmentation or predefined cell annotations. We found that other models performed unsatisfactorily, with ROC-AUC values near 0.5 (Fig. 4f). In contrast, TissueFormer achieved a mean ROC-AUC of 0.684, with statistically significant gains (*P <* 0.01) over all baselines (Fig. 4f). Remarkably, despite relying solely on H&E images, TissueFormer performed on par with an oracle variant (TissueFormer-Oracle) that uses measured gene expression as input on 4 out of 12 cases. The spatial niche predictions produced by TissueFormer also aligned with biological expectations: in tissue samples with increasing fibrosis severity, we observed a reduction in the healthy alveolar niche (T4) and an increase in diseased niches (T6, T9) (Fig. 4g, fig. S14). Moreover, we compared our model with recent specialized spatial transcriptomic models, Streamboat [43], MintFlow [44] and iSCALE [45], in the spatial niche lassification task. All three models use as input spatial transcriptomic data but no histology images. For a fair comparison with the embedding learned by these models, we used only TissueFormer’s embedding from the spatial transcriptomic data. The embeddings served as input for a three-layer MLP which is trained to classify cells into spatial niches. The results show that TissueFormer significantly outperforms these recently proposed models across the 12 different cases, supporting the utility of our model (fig. S15).

Overall, these results demonstrate that TissueFormer delivers accurate and biologically grounded predictions across whole-slide, region, and single-cell levels from histology images. It establishes a unified framework that bridges routine histopathology with molecular and diagnostic insights.

### Attention mechanism reveals long-range cellular dependencies from histology images

The attention mechanism in TissueFormer enables quantification of dependencies between arbitrary pairs of cells across an entire tissue slide, offering a powerful tool for *in silico* analysis of cellular dependencies at scale. Unlike conventional spatial analyses that rely primarily on local adjacency, TissueFormer captures long-range dependencies often overlooked yet critical for understanding tissue organization, disease progression, and therapeutic response [30–32]. For each slide, we input the H&E image into TissueFormer ‘s histology encoder, whose graph transformer outputs an all-pair attention matrix. This matrix assigns a pairwise attention score (ranging from 0 to 1) that reflects the learned influence between any two cells, regardless of spatial proximity (Methods), producing a data-driven, fine-grained map of cell–cell dependencies. This cell-level, attention-based analysis—particularly capturing long-range dependencies in high-resolution tissue images—is uniquely enabled by TissueFormer through its scalable global attention mechanism.

As a first case study, we applied TissueFormer to tissue slides from the same pulmonary fibrosis (PF) patient dataset [36], analyzing how intercellular attention patterns evolve with disease progression. PF is a progressive, lethal interstitial lung disease characterized by aberrant fibroblast proliferation and extracellular matrix deposition, leading to irreversible alveolar damage and respiratory failure. Airspace regions–the primary sites of gas exchange—-are especially vulnerable to injury, and their deformation strongly correlates with mortality in PF patients [15, 36]. We analyzed attention levels between activated fibrotic fibroblasts—key effector cells in PF—and immune cells with disparate spatial proximity (within or outside ROI), across ROI’s pseudo time, a measure of disease severity from healthy to late-stage fibrosis (Fig. 5a, Methods). We observed that attention levels from activated fibrotic fibroblasts to immune cells in airspace regions increased with disease progression, regardless of spatial proximity (Fig. 5b). However, the time point of this increase varied across cell types and spatial distances. Typically, attention levels on alveolar and SPP1^+^ macrophages in airspace regions rose markedly at late stages, while attention levels on the same macrophage types outside airspace regions increased at earlier stages (Fig. 5b–d; fig. S16, fig. S17). This indicates long-range fibroblast–macrophage affinity occurring earlier in PF progression.

**Fig. 5.**
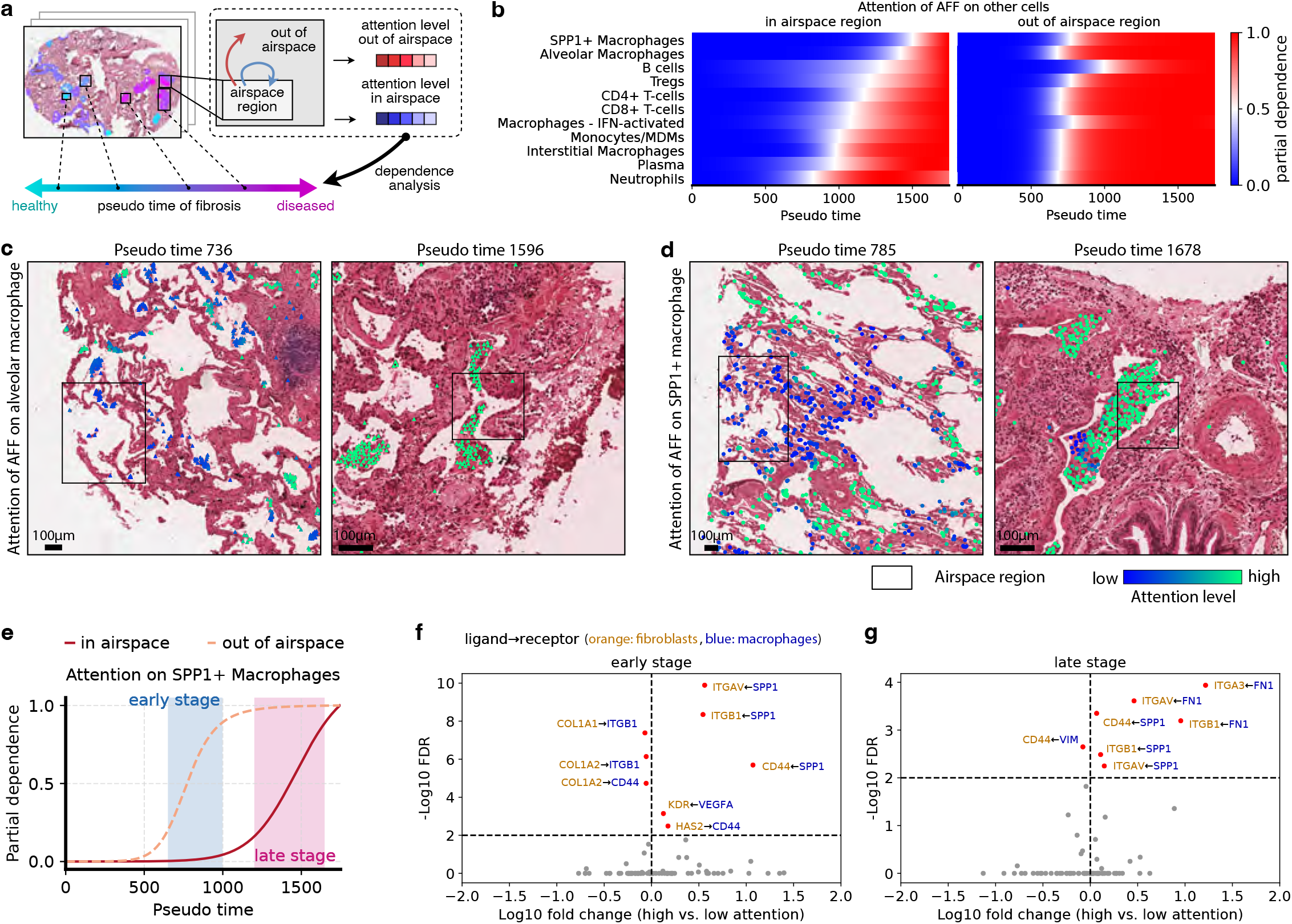
Attention-based modeling and analysis of cell-cell dependencies. **a**, Illustration of the analysis on tissue slides with pulmonary fibrosis, where we utilized pretrained TissueFormer to output pairwise attention levels between cells of each tissue slide. We investigated the attention between cells inside the airspace and the attention of cells inside the airspace on those outside of it, and further analyzed their dependence with airspace space’s pseudo time, a reflection of the disease severity of airspace regions. **b**, Plots showing partial dependence of attention levels of activated fibrotic fibroblasts (AFF) in the airspace on immune cells (with significance level *P <* 0.01) in and out of airspace, respectively, with respect to pseudo time. **c**, Representative examples from two pseudo times that correspond to the increasing points of attention levels on alveolar macrophages out of (left image) and in the airspace (right image). Overlaid on the H&E image we show the alveolar macrophages and the colors indicate the attention levels of activated fibrotic fibroblasts on alveolar macrophages. **d**, Representative examples for SPP1^+^ macrophages from two pseudo time points. **e**, Plot showing the partial dependence of attention levels of activated fibrotic fibroblasts on SPP1^+^ macrophages in and out of the airspace, respectively. The early stage and late stage are marked by the increasing intervals of attention levels. **f**, Differential analysis for co-expression levels of ligand-receptor pairs from cell pairs (each pair consists of an activated fibrotic fibroblast and a SPP1^+^ macrophage) with high (above 80 percentile) and low (below 20 percentile) attention levels. The fold change is the ratio of the averaged co-expression level from the high-attention group to that of the low-attention one, and FDR *>* 0.01 is set as the threshold of statistical significance.

Focusing on SPP1^+^ macrophages (referred to as macrophages below), a more homogeneous population than alveolar macrophages, and activated fibrotic fibroblasts (referred to as fibroblasts below), we examined the biological meaning of the attention scores (fig. S18a). We found that the cellular attention maps produced by TissueFormer are strongly associated (p ¡ 0.001) with gene expression predictability. Specifically, the image embedding of a cell receiving high attention (e.g., a macrophage) from a querying cell (e.g., a fibroblast) exhibits a greater capacity to predict the querying cell’s gene expression profile than embeddings of cells receiving low attention. Interestingly, this analysis reveals that distal cells—located well beyond the typical range of intercellular communication—can still be predictive of the state of a cell of interest. The long-range interctions reflect latent semantic relationships between non-adjacent cells with related morphological and molecular features. To further assess the biological relevance of the learned attention maps, we examined ligand–receptor (LR) co-expression between fibroblasts (querying cells) and macrophages (cells receiving high attention); see Fig. 5e–g; fig. S19. We found that at early disease stages, fibroblast-macrophage attention levels were significantly associated with the co-expression of the ligand SPP1 (expressed by macrophages) and its common receptors, integrins and CD44 (expressed by both macrophages and fibroblasts), suggesting possible recruiting signals from fibroblast to macrophages. By contrast, in late-stage fibrosis, these signals waned and the attention levels exhibited strong association with the co-expression of the pro-fibrotic gene Fibronectin1 (FN1) (expressed by macrophages) and common SPP1-receptors (expressed by fibroblasts). This is in line with previous studies recognizing that high SPP1 expression converts macrophages to a pro-fibrotic M2-like phenotype [46], and that a population of pro-fibrotic macrophages marked by expression of FN1 expands following organ injury (i.e., late-stage fibrosis) [47]. To further support the biological relevance of the learned attention maps, we performed differential gene expression analysis between macrophages receiving high versus low attention, stratified by early and late stages of disease (fig. S18c–e). In early-stage disease, macrophages receiving high attention exhibited increased expression of chemotactic and infiltrative signaling genes compared to those receiving low attention (e.g., *CCL18, CXCR4, FCN1*). In contrast, in later stages of disease, the differences were primarily observed in metabolic and stress-response genes, including *SLC2A1* (GLUT1), *HIF1A*, and *HSPA5* (BiP). These results unbiasedly reveal a dynamic fibroblast–macrophage dependency across disease progression, with macrophages transitioning from signaling hubs in the interstitium to metabolically stressed, matrix-remodeling cells within fibrotic regions. These findings are consistent with prior studies showing that SPP1^+^ macrophages are enriched for secreted factors and inflammatory/immune pathways that promote fibroblast activation [48], and that elevated SPP1 expression drives macrophages toward a pro-fibrotic, M2-like phenotype in late-stage organ fibrosis [46, 47].

As a second case study, we applied TissueFormer in the context of breast cancer to tissue sections from patients with invasive cancer and Ductal Carcinoma in Situ (DCIS), an early form of cancer further categorized into different stages (DCIS1, DCSI2) [37]. Our goal was to explore whether intercellular attention patterns could enhance cell subtype identification beyond what is achievable using intrinsic molecular features alone (fig. S20a). To characterize immune–tumor cellular dependencies in the cancer microenvironment, we aggregated attention scores by averaging the attention received by each cell type. This revealed distinct patterns of immune–tumor crosstalk consistent with prior knowledge (fig. S20b–c). For instance, both DCIS1 and DCIS2 exhibited strong affinity with macrophages, while invasive tumor cells engaged more frequently with CD4^+^ T cells (fig. S20c), consistent with previous studies reporting a role of macrophages in breast cancer early dissemination [49], and of T cells in later more invasive stages [50, 51]. To dissect tumor heterogeneity further, we analyzed subtype-level interactions, focusing on how macrophage–tumor dependency patterns stratify cancer cells into more refined subpopulations (fig. S20d–f). Using macrophage–DCIS1 attention patterns, we identified two subtypes, both of which displayed spatial continuity. Applying the same method to DCIS2 yielded two additional subtypes. Together, these four subtypes (DCIS1-a, DCIS1-b, DCIS2-a, DCIS2-b), which could not be identified based on gene expression analysis alone, formed a malignancy gradient (fig. S20f), with DCIS2-c appearing most aggressive and DCIS1-b most benign—offering a more fine-grained stratification than clustering based on intrinsic features alone. Conversely, examining the attention of tumor cells on macrophages revealed functionally distinct subtypes of macrophages (fig. S20g-j). For DCIS1, two macrophage subtypes emerged: one displaying regulatory, tissue-resident (M2-like) profiles (e.g., *CX3CR1, CD14, PPARG, CD80*), and the other marked by phagocytic and chemotactic signatures (e.g., *ITGAX, CD68, FOGR1A*) (fig. S20g-h). For DCIS2, macrophage subtype 2 exhibited an inflammatory M2-like state (*LYZ, CD14, CD163, FCER1A*), whereas macrophage subtype 1 was enriched for antigen-presenting markers (*CD80, FCGR3A*) (fig. S20i-j). Overall, we found that macrophages are the most predictive cell type of DCIS, and that distinct macrophage subtypes exhibit co-dependencies with DCIS subtypes of varying malignancy grades.

Together, these findings highlight TissueFormer’s attention-based mechanism as a powerful computational tool for intercellular communication analysis. Its interpretable, high-resolution spatial attention maps can be used to reveal long-range cellular dependencies and enable refined characterization of cancer cell subtypes, paving the way for scalable spatial discovery and disease understanding at single-cell resolution.

## 3 Discussion

We have presented TissueFormer, a multimodal foundation model pretrained on a large corpus of whole-slide histology images paired with spatial transcriptomic profiles. TissueFormer achieves strong performance in predicting molecular profiles and diagnostic annotations directly from routinely acquired histology images, and generalizes effectively across tissues, species, and a variety of downstream tasks. Technically, TissueFormer addresses two fundamental challenges in spatial modeling at single-cell resolution. First, it introduces a whole-slide graph transformer that captures both local dependencies among adjacent cells and global, long-range dependencies across cells in a slide. This is enabled by a scalable attention mechanism that computes all-pair interactions with linear complexity in the number of cells. Second, TissueFormer employs a scalable multimodal contrastive loss that facilitates efficient training by aligning image- and expression-based representations through all-pair contrasts across cells, while similarly reducing computational complexity from quadratic to linear. These innovations provide the scalability necessary to train on high-resolution slides with up to millions of cells each. Furthermore, while we specifically focused on modeling cells in space, our technical innovations could significantly reduce computational costs and benefit any foundation model where long-range dependence modeling is important. This includes, for example, models that capture interactions between amino acids to define protein folding, or those that capture interactions among distant nucleotides that influence gene expression.

Our evaluation results demonstrate that TissueFormer can be deployed for prediction tasks on new datasets without retraining the model. Using an in-context learning approach, TissueFormer achieves competitive performance by referencing only a small number of slides from the target dataset—an especially valuable capability given the high cost of spatial transcriptomics. This allows TissueFormer to serve as a practical and scalable framework for “virtual spatial transcriptomics”, reducing experimental burden by computationally inferring spatial gene expression from standard histology images with minimal paired data.

In addition to its predictive power, TissueFormer offers a new computational tool for *in silico* cell–cell interaction analysis. Traditional approaches typically restrict analysis to locally adjacent cells, relying on fixed-radius thresholds or nearest-neighbor heuristics. These methods cannot identify long-range intercellular dependencies and require manual parameter tuning. In contrast, the global attention mechanism in TissueFormer enables fully data-driven inference of interaction between any pair of cells in a slide, regardless of spatial proximity. This affords greater flexibility and expressivity for analyzing tissue organization and cell-cell signaling, with direct implications for understanding disease mechanisms and identifying long-range cellular dependencies.

Potential future directions include applying TissueFormer to gene panel design, where gene expression predicted from histology images can guide the selection of a targeted subset of genes relevant to specific diseases, thereby reducing the experimental costs for biomarker identification. Additionally, the attention-based cell–cell interaction analysis could be extended to incorporate gene perturbation data with spatial resolution, to model the effects of disease-associated perturbations on cellular dynamics, potentially enhancing the identification of therapeutic targets associated with early-stage disease.

## 4 Methods

### 4.1 Data preprocessing

We employed the following preprocessing pipeline for multi-modal spatial data including histopathology (H&E-stained) images, gene expression from spatial transcriptomics profiles, and cell-cell spatial adjacency.

#### Histology images

For H&E-stained images, following Prov-GigaPath [24], we rescaled each image slide to a standard resolution of 0.5 *µ*m per pixel, i.e., 20 × magnification. This step is to make sure that all image slides as input of the model have a unified spatial resolution. Then for each cell of a Xenium slide (respectively spot of Visium, Visium HD or ST slide), we extracted a 224 × 224-pixel image patch centered around each cell nucleus (respectively spot). This yielded 1,593,892 spot-centered and 15,434,350 cell-centered image patches in total from the HEST-1K dataset used for pretraining.

#### Gene expression profiles

For spatial transcriptomic profiles, we first unified gene names by filtering special symbols and linking genes with different synonyms [52]. Since different tissue slides can have largely disparate measured gene panels, we identified a union of genes from all tissue slides which consists of about 20K human protein-coding genes. We filtered transcripts with QV above 20. For Xenium slides, transcripts within each nucleus were aggregated to obtain a cell-by-gene matrix. For Visium, Visium HD or ST slides, transcripts within each spot were aggregated to obtain a spot-by-gene matrix. Following common practice, we normalized expression values by total count per spot or cell and did log1p transformation. For evaluation purposes, we calculated the highly variable genes (HVG) for each tissue slide with Seurat v3 [53].

#### Spatial adjacency graph

Besides histology and expression features, we constructed a cell-cell (or spot-spot) adjacency graph. For each tissue slide, we used the spatial locations of cells or spots in the coordinate system to compute their pairwise Euclidean distances, and the adjacency graph was constructed by connecting each cell or spot with its five nearest neighbors. This yielded averaged 14,190 nodes and 70,950 edges per tissue slide graph, the largest graph in the dataset containing 1,355,603 nodes and 6,778,015 edges.

### 4.2 Model architecture

The TissueFormer model consists of a histology encoder and an expression encoder. The histology encoder takes a whole-slide H&E-stained image as input and outputs the embeddings of each single cell. Each cell is represented by a 224 × 224-pixel cell-centered image patch which is fed into an encoder [35] pretrained with over one million H&E images. This converts each cell-centered image patch into a 1536-dimensional dense vector as a cell-level embedding. These cell-level image embeddings together with the spatial adjacency graph are then fed into our *whole-slide graph transformer* model to compute context-aware image-based cell representations.

#### Whole-slide graph transformer

For a tissue slide with cells indexed by *i* ∈ {1, · · · , *N* } = N, denote by **x**_*i*_ the image embedding for cell *i*. Besides, denote by **A** = [*a*_*ij*_]_*i,j*∈N_ the cell-cell adjacency matrix, where *a*_*ij*_ = 1 if cell *i* and *j* are connected in the adjacency graph and *a*_*ij*_ = 0 otherwise, and *d*_*i*_ denotes the degree of cell *i* in the graph, i.e., *d*_*i*_ = ∑_*j*∈N_ *a*_*ij*_. The transformer model first maps input image embeddings into initial embeddings 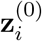 by a shallow multi-layer perceptron (MLP):

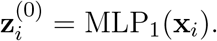

Then the transformer model updates the cell embeddings via a stack of *L* propagation layers. Namely, the *k*-th propagation layer updates cell embeddings from 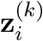 to 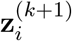 via the updating rule:

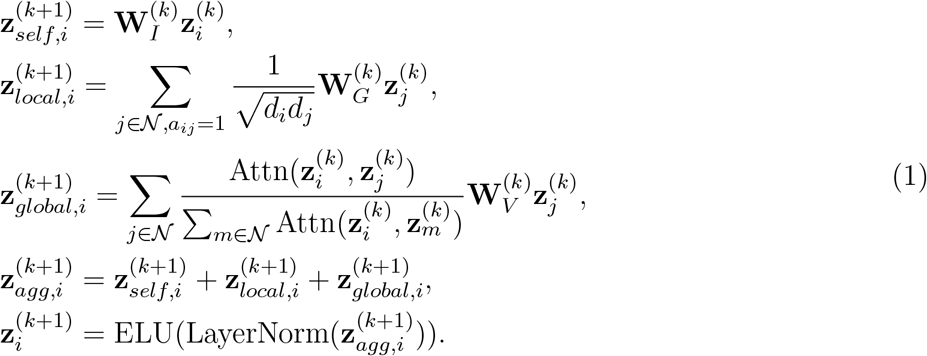

The first term 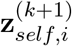, serving as self-loop propagation, models the intrinsic effect of each cell itself. The second term 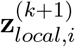, which inherits the propagation layer of the graph convolutional network (GCN) [54], captures the local effect of neighboring cells. The third term 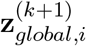 implements a whole-slide attention mechanism that accommodates the global effect of all cells in a tissue slide. Then the three-fold effects are aggregated together in 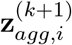 to compute the next-layer embeddings 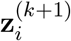, where ELU and LayerNorm denote ELU activation and layer normalization, respectively. In terms of the GCN, our exploratory analysis suggests that using relatively small neighborhood sizes for constructing the spatial adjacency graph enables the optimal performance (fig. S21a). The key design of the transformer model lies in the whole-slide attention mechanism, which, however, introduces a non-trivial computational obstacle, since the attention computation requires *O*(*N* ^2^), i.e., quadratic complexity with respect to the number of cells, for computing all 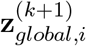, *i* ∈ *N* . We overcome this challenge by introducing a special attention mechanism that reduces the computational complexity to linear order *O*(*N*) while accommodating dependencies among arbitrary pairs of cells.

#### Scalable whole-slide attention mechanism

We specify the attention network Attn(·, ·) as a dot-product function in unit sphere space:

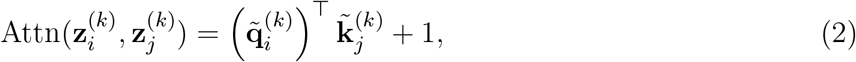

where the query and key vectors are normalized by their L2 norms, respectively:

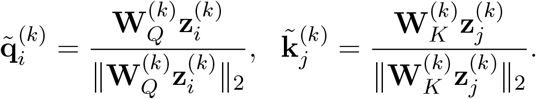

Here 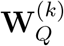 and 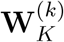 are trainable weight matrices of the *k*-th layer. In Eqn. (2), the addition of 1 to the dot-product in unit sphere space guarantees the non-negativity of the attention result. While directly computing the standardly used global attention term 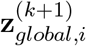 in Eqn. (1) is highly time- and space-consuming (with *O*(*N* ^2^) complexity) making it infeasible for training at single-cell resolution at whole-slide scale, our proposed attention mechanism enables an efficient computation scheme as described below.

Assume 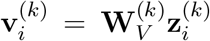 to simplify the notation. We notice the following equivalent matrix multiplication according to the association rule in linear algebra:

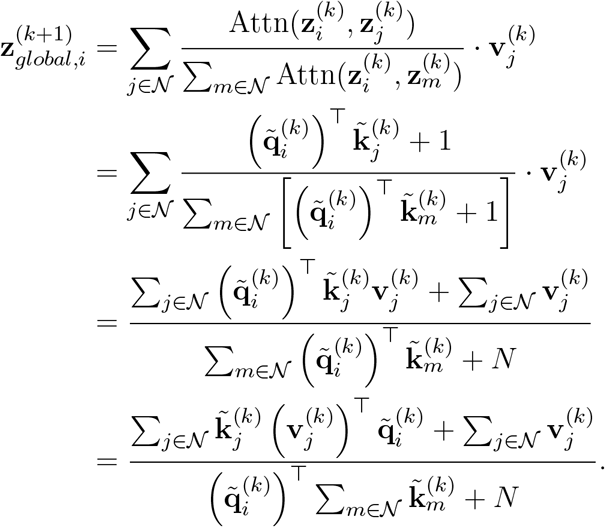

Since the summation terms in the right-hand side are shared across all cells *i* ∈ N, we only need to compute them once, meaning that the total complexity for computing all 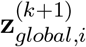, *i* ∈ N, can be achieved within *O*(*N*), i.e., linear-order complexity with respect to the number of cells *N* . To understand how we achieve computation of this all-pair attention within *O*(*N*) complexity in implementation, we can use the matrix form for a further explanation. Denote by 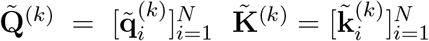 and 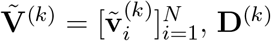 is an *N* × *N* diagonal matrix for normalization (from the attention’s denominator) and **I** is an *N* × *N* identify matrix. Then the updated node embeddings from *k*-th to (*k* + 1)-th layer would be:

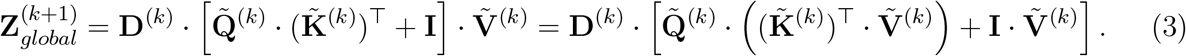

Derivation from the left-hand-side to the right-hand-side (which is adopted for implementation) does not result in changes of results but reduces the computational complexity from *O*(*N* ^2^) to *O*(*N*).

Notice that the above acceleration does not require any approximation, so the global attention between any pair of cells is still fully accommodated, making it feasible for scaling up to high-resolution image slides without compromising the model’s expressivity. The image-based cell representations, denoted by **r**_*i*_, are computed by stacking multiple whole-slide graph transformer layers. Our exploratory analysis suggests that using a shallow transformer architecture is sufficient to produce practically relevant efficacy levels (fig. S21b).

#### Expression encoder

For the gene expression encoder, we first map each gene into an embedding in latent space. Denote by G the union set of genes (obtained by preprocessing) for pretraining, and we consider a trainable look-up table of gene embeddings **E** = {**e**_*k*_}_*k*∈G_. For a tissue slide with gene panel P ⊂ *G*, denote by **s**_*i*_ = [*s*_*ik*_]_*k*∈P_ the gene expression vector of cell *i* where *s*_*ik*_ is the expression value of gene *k*, and we compute the cell embedding by aggregating the gene embeddings:

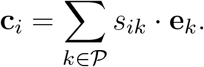

The gene expression-based cell representations **b**_*i*_ are then computed through another shallow MLP:

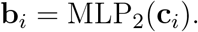

To keep the notation clean, we denote by *f*_1_ the histology encoder that gives rise to the image-based cell representations **r**_*i*_ = *f*_1_(**x**_*i*_) and by *f*_2_ the expression encoder that outputs the gene expression-based cell representations **b**_*i*_ = *f*_2_(**s**_*i*_).

### 4.3 Pretraining

The pretraining of TissueFormer resorts to contrastive alignment between the histology encoder *f*_1_ and the expression encoder *f*_2_ to take advantage of the paired modalities in the training data. To this end, we design a multi-modal contrastive learning objective that exploits self-supervision through bi-directional learning between the image representations and gene expression representations that are considered as two views of each cell.

#### Multi-modal contrastive learning

Our contrastive learning approach treats the two-view representations of the same cell as a positive pair and aims at minimizing their discrepancy, while it defines the representations of different cells as negative pairs and maximizes their differences. From an information-theoretic viewpoint, the objective can be interpreted as a variational lower bound of the mutual information between two-view representations I(*f*_1_(**x**_*i*_, *f*_2_(**s**_*i*_)) [55]. With the commonly used L2 distance metric ∥· ∥ _2_ for alignment between two-view representations, the symmetric version of the multi-modal contrastive learning loss can be written as [56–58]:

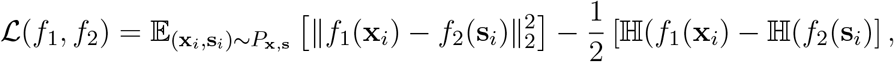

where E denotes the expectation taken over *P*_**x**,**s**_, the joint distribution of the input data, and H denotes the entropy of the representations. The two entropy regularization terms that essentially maximize the discrepancy between representations of different cells disincentivizes a degenerate solution where two encoders map to a constant [59]. Inserting **r**_*i*_ = *f*_1_(**x**_*i*_) and **b**_*i*_ = *f*_2_(**s**_*i*_) into the objective leads to the following contrastive learning loss for one tissue slide:

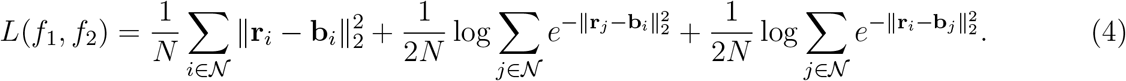

Note that the two entropy terms require computation of the distances between any pair of cells, inducing *O*(*N* ^2^) complexity, which again hinders scalability to whole-slide training at single-cell resolution. While a straightforward solution is to sample a mini-batch of cells as negative pairs, this would inevitably introduce sampling bias and lead to inaccurate estimation of the target objective. To overcome this problem, we introduce a new computational scheme for the above contrastive learning loss that reduces the computational complexity to *O*(*N*) without the requirement of mini-batch sampling.

#### Scalable contrastive loss

We first map the representations to a unit sphere:

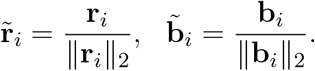

In this way, the pairwise distance is equivalent to the negative of the dot-product similarity:

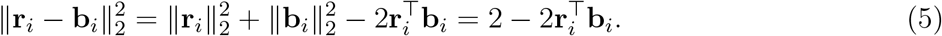

Furthermore, by using the first-order Taylor expansion to approximate the exponential function, i.e., *e*(**x**) ≈ 1 + **x** for any small **x** ∈ R, the loss function to be minimized can be derived from Eqn. (4) as

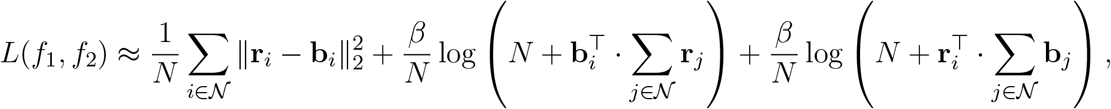

where *β* is the weight for the entropy regularization, which we set to 0.5 by default. Similar to the computation of the global attention, both summation terms inside the logarithm are shared across all cells *i* ∈ *N*, so they only need to be computed once in practice. This reduces the time and space complexity of computing *L*(*f*_1_, *f*_2_), the contrast between any pair of cells in a whole tissue slide, to *O*(*N*).

#### Training details

We used Xavier uniform initialization for the gene expression embeddings **E** and the default initialization in PyTorch for the other model layers. For each training iteration, we feed one whole tissue slide into the model for the feed-forward computation and back propagation, which can be accomplished on a standard GPU thanks to our scalable computation schemes. To facilitate convergence and generalization, we utilized a two-stage curriculum training strategy. For first-stage training, we trained the model from scratch on low-resolution data (sequencing-based slides with spot-level gene expression), with learning rate 10^−5^ for 1000 epochs. To stabilize the training over large heterogeneous data, we updated the model parameters with the accumulated gradients of 20 iteration steps. Then for second-stage training, we fixed the gene expression embeddings and trained the model on high-resolution data (Xenium slides with cell-level gene expression), with learning rate 10^−5^ for 200 epochs. The model pretraining utilized 8 × 50 GB A6000 GPU and took approximately two days. The inference time on a whole tissue slide was on average 0.4 seconds.

### 4.4 Inference and finetuning

After pretraining, the TissueFormer model can be applied to multiple diverse tasks related to cross-modality retrieval, multi-level prediction and attention interpretation. All of the tasks demonstrated in this paper are based on histopathology (H&E) images as inputs. For any test tissue slide that has a whole-slide H&E image as observation, with cells indexed by *i* ∈ {1,· · ·,*N* ^′^} = *N*^′^, we first compute the cell-level image patch embeddings 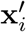 and cell-cell adjacency matrix. They are then fed into the pretrained histology encoder *f*_1_ of TissueFormer to obtain image-based cell representations denoted by 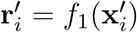 for cell *i* ∈ N^′^.

#### Cross-modality retrieval

As the histology encoder was pretrained through alignment with the paired gene expression profiles, the image-based cell representations enable predicting gene expression for test tissue slides that only have H&E images. To this end, inspired by recent advances in large language models with zero-shot generalization capabilities [20, 60, 61], we propose a training-free in-context learning approach. Specifically, we use the paired samples (that can be a subset of the training data or additional data) denoted by 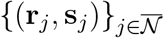, where 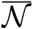 denotes the union of cell indices from training tissue slides, as *reference*. Then for any cell *I* ∈ *N*^′^ from a test tissue slide, we treat its image-based cell representation 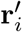 as a *prompt* and retrieve the top *K* cells in 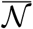 that are closest to 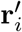 in latent space:

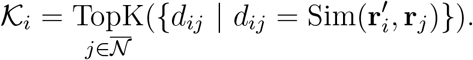

We specify Sim(·, ·) as the dot-product function after L2 normalization of the input representations. Then the predicted gene expression vector ŝ_*i*_ as an *answer* is estimated through a weighted sum of those of the retrieved cells:

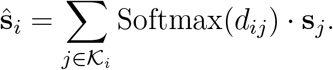

The gene expression vector **s**_*j*_ can be flexibly specified or extended with gene markers unseen in training, which enables the model for zero-shot generalization to new unseen genes without any retraining.

#### Multi-level prediction

Apart from predicting gene expression, the pretrained TissueFormer model can be used to predict cell-level labels *ŷ*_*i*_. To achieve this goal, a task-specific linear probing layer or shallow MLP is trained as the *predictor* that uses the image-based cell representations 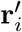 and the predicted gene expression-based representations 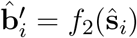:

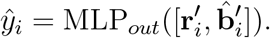

We train the predictor with MSE loss for regression tasks and the Focal loss [62] that works smoothly for classification tasks with imbalanced labels. For handling region-level tasks, we use sum pooling over the representations of cells within a region of interest (ROI) to obtain region-level representations, which are then fed into the predictor. Similarly, for slide-level tasks, we take sum pooling over cell representations of a whole slide to obtain slide-level representations.

#### Attention interpretation

The training of TissueFormer does not explicitly compute the pairwise attention between cells, as our scalable scheme avoids the appearance of the *N* × *N* all-pair attention matrix in the feed-forward computation. At test time, however, we can extract the attention for any pair of cells from the pretrained model. Specifically, with the cell-level image patch embeddings **x**_*i*_ as input, we use the pretrained histology encoder of TissueFormer to compute the unnormalized attention score *o*_*ij*_ between cell *i* and cell *j* via the trained attention network Attn(·, ·). In this way, we can obtain a whole-slide all-pair attention matrix [*o*_*ij*_]_*i,j*∈N_ , based on which we extract cell-specific and cell-type-specific attention levels for further analysis.

### 4.5 Evaluation of Gene Expression Prediction

For the gene expression prediction task, we first evaluated the model with different splits of tissue slides from HEST-1K [34] to investigate the generalization capability of the model. We used the Visium, Visium HD and ST slides (1136 in total) that give rise to a shared gene panel of about 4K human protein-coding genes, and considered two data splits for evaluation which divided tissue slides with different organs and species, respectively. In the setting of generalization across organs, we split the 21 organs into “frequent” organs (i.e., with a lot of data), namely the top five organs (spinal cord, brain, breast, bowel and skin) in terms of number of slides, and the other organs (lung, kidney, heart, prostate, liver, uterus, muscle, eye, bone, pancreas, bladder, lymph node, cervix, ovary, embryo and placenta). We used 702 slides of the frequent organs for training, and tested the model on 100 held-out slides of these five organs as well as 334 slides of the sixteen unseen organs. In the setting of generalization across species, we trained the model on slides of mus musculus and tested on held-out slides of homo sapiens, as compared with the counterpart trained on slides of homo sapiens and tested on held-out slides of homo sapiens (in the two scenarios the same slides were used for testing).

For benchmarking, we compared with state-of-the-art pretrained foundation models (H-optimus-1, Prov-GigaPath and UNI) and a non-deep-learning method PCA as competing models. Akin to TissueFormer, the three foundation models only used H&E images of test tissue slides as input and output spot-level representations, while the PCA calculated the top 50 principle components of H&E images as spot-level representations. The spot-level representations were then fed into a linear regression layer to predict spot-level gene expression. The linear regression layer was trained with the training data as described above, with 100 epochs and learning rate 10^−5^. Performance was measured by Spearman correlation which was first calculated per spot between the predicted and ground-truth gene expression of the top *n* HVG (*n* = 4000, 2000, 1000) and then averaged across spots to obtain a slide-level score.

We also investigated the model’s prediction accuracy for high-resolution gene expression at cellular level. We trained the model through curriculum learning on all Visium, Visium HD and ST slides as well as Xenium slides except the ones held-out for testing. The test Xenium slides from HEST-1K (TENX126, TENX124, TENX123, TENX121, TENX119 and TENX118) come from six different organs (pancreas, lymphoid, skin, liver, heart and lung) but share a unified gene panel, which allowed for evaluating the model’s prediction on specific gene markers across different organs. The performance was measured by gene-wise Mean Squared Error (MSE) that averages the square distance between predicted and ground-truth expression of a given gene across cells.

To evaluate the model’s capability for generalization to new gene markers unseen in training, we adopted three bone tissue slides (TENX135, TENX136 and TENX137) from HEST-1K that were used for training, and tested the model on 40 genes out of their gene panel that were randomly held out from training. We adopted slide-level cross-validation for evaluation; namely, we used any two of three tissue slides as reference for in-context learning and predicted gene expression on the remaining slide. We reported the MSE of every unseen gene. The conditional entropy ℍ (*k*^′^|G) between a given unseen gene *k*^′^ and the union of training genes G was calculated via:

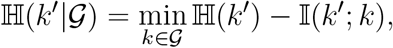

where the entropy H(*k*^′^) and the mutual information I(*k*^′^; *k*) were estimated using gene expression histograms.

To further validate the model’s capability to generalize to new datasets, we used lung tissue slides with pulmonary fibrosis [36] as an independent dataset for evaluation. We randomly held out 20 tissue slides from 36 in total for testing. To comprehensively investigate the model’s generalization capability, we compared the predictive performance in two settings: 1) zero-shot generalization that used tissue slides from HEST-1K as reference for prediction; 2) few-shot generalization that used different numbers of (1-16) tissue slides of the evaluation dataset as reference. Performance was measured by Spearman correlation between predicted and ground-truth cell-level gene expression averaged across all cells of the testing slides. Moreover, we evaluated the prediction on gene markers associated with pulmonary fibrosis. These gene markers were identified via fitting a linear model between gene expression and percent pathology scores [36], filtering out the genes which were not statistically significant (FDR ≥ 0.01) on the test slides, leading to 40 positively disease-associated markers and 28 negatively associated ones. We compared Pearson correlation between ground-truth gene expression and pathology scores of the test slides with the Pearson correlation between the predicted gene expression and pathology scores of the test slides.

### 4.6 Evaluation on digital diagnosis tasks

Beyond predicting gene expression, we applied the TissueFormer model pretrained on all tissue slides from HEST-1K to various prediction tasks on the pulmonary fibrosis dataset [36] to demonstrate the wide capabilities of TissueFormer for informing digital diagnosis at different biological scales. The tasks of interest included patient stratification, ROI disease severity inference, disease-associated cell type detection, pathology feature annotation and spatial niche classification. These tasks are all based on H&E images as input and can be categorized into slide-level, region-level and cell-level tasks.

Following the previous data splits, we used 20 tissue slides for testing and the remaining 16 slides for training. The H&E images and gene expression profiles of the training slides were used to finetune the TissueFormer model with the contrastive learning loss. For the test slides, we used H&E images to obtain image-based cell representations as well as predicted gene expression that were used to obtain predicted gene expression-based cell representations. These representations were used for handling different tasks.

For patient stratification, we aggregated the image-based cell representations to compute slide-level representations via sum pooling, which we visualized using UMAP.

For ROI disease severity inference, airspace regions of tissue slides were considered as regions of interest (ROI) and the goal is to predict the disease severity labels of ROI. We trained a shallow MLP as the predictor based on the region-level representations, which were obtained by sum pooling over the gene expression-based cell representations, to predict disease severity labels. We used MSE loss and trained the predictor with 400 epochs and learning rate 10^−4^ on the training slides. For inference on test slides, we aggregated the predicted gene expression-based representations over the cells of each ROI to obtain region-level representations and used the trained predictor for prediction. Then we compared the ground-truth and predicted disease severity labels of the test slides.

For disease-associated cell detection, we trained an MLP classifier to predict from the gene expression-based cell representations the four cell types (capillary, transitional AT2, activated fibrotic fibroblasts and SPP1^+^ macrophages) that mark the transition of disease states of fibrosis. We trained the predictor with 100 epochs and learning rate 10^−5^ using Focal loss (*α* = 0.25, *γ* = 2) for binary classification, where we randomly sampled five negative samples per positive sample. For inference on test slides, we fed the predicted gene expression-based cell representations into the predictor for prediction. To evaluate whether the predicted cell types preserve the evolution trends of ground-truth cell ratios within ROI, we compared the normalized cell ratios (ratio of a cell type normalized between 0 and 1) in ROI to those from the prediction. For pathology feature annotation, we trained an MLP predictor that used image-based cell representations to predict ten pathology features (severe fibrosis, epithelial detachment, fibroblastic focus, hyperplastic AECs, large airway, giant cell, granuloma, advanced remodeling, small airway and venule). The predictor similarly trained with Focal loss using a 1:5 ratio for negative sampling. For benchmarking, we compared with H-optimus-1, Prov-GigaPath and UNI as competing methods, and for these models, we also trained the MLP predictor using their image-based cell representations. For model selection, we searched over learning rates in {10^−2^, 10^−3^, 10^−4^, 10^−5^, 10^−6^, 10^−7^} and training epochs in {100, 10, 1} for each model on the validation set held out from the training slides. Predictive performance was measured using ROC-AUC scores.

For spatial niche classification, the goal is to classify cells into spatial niches. We used the transcript-based spatial niche labels provided by the original paper [36] that were agnostic to cell segmentation and comprehensively defined spatially integrated molecular units relevant to the lung disease. The original paper labeled these spatial niches with T1-T12, and we named them according to the cell type composition provided by [36] to facilitate the interpretation of the results. For TissueFormer, we trained an MLP to predict from the gene expression-based cell representations the spatial niche labels. At inference time, the predicted gene expression was used as input for prediction on the test slides, while for TissueFormer-Oracle, we used the ground-truth gene expression. For the competing methods, the image-based cell representations were used to train the MLP predictor for classification. Similar to the pathology feature annotation task, we searched over learning rates in {10^−2^, 10^−3^, 10^−4^, 10^−5^, 10^−6^, 10^−7^} and training epochs in {100, 50, 30} for model selection. We reported the ROC-AUC scores for each niche label.

### 4.7 Attention-based analysis

The pretrained TissueFormer model enables the extraction of an intercelluar attention matrix **O** = [*o*_*ij*_]_*i,j*∈N_ for each tissue slide. This attention matrix quantifies the pairwise dependence between any pair of cells in the whole slide, where high (low) attention scores indicate that the two cells tend to have strong dependencies.

For analysis in the lung fibrosis case, we extracted a submatrix from the whole-slide attention matrix by constraining to cells around each airspace region. In this way, we obtained an attention submatrix **O**_*m*_ that reflects the cellular dependencies within and near ROI *m*. Based on this, we calculated a cell-type-specific attention vector **p**_*m*_ by averaging the attention scores over each cell type. Specifically, we investigated the attention levels from activated fibrotic fibroblasts to immune cells, denoted by **p**_*m*_, and then used a generalized additive model (n spline = 5, lam = 1) to fit **p**_*m*_ with the pseudo time label *t*_*m*_ of ROI *m*. Based on the results, we zoomed in on SPP1^+^ macrophages whose attention levels from activated fibrotic fibroblasts exhibited significant difference depending on spatial proximity: the attention levels on macrophages outside of airspace regions increased at the early stage, while the attention levels on macrophages inside airspace regions increased at the end stage. We next analyzed the ligand-receptor (LR) pairs that are highly correlated with the increasing patterns of the attention levels. For this purpose, we extracted all cell pairs containing activated fibrotic fibroblasts and SPP1^+^ macrophages from a given ROI, and divided these cell pairs into a high-attention group (with attention levels above the 80th percentile) and a low-attention group (with attention levels below the 20th percentile). For each LR pair, we computed co-expression levels of the two genes (multiplication of cell-level expression of two genes) per cell pair, and then calculated the fold-change of co-expression levels between the high- and low-attention groups. Specifically, we measured the fold-change by the ratio of the mean co-expression level from the high-attention group over that of the low-attention group.

For the analysis in the breast cancer case, we extracted the attention matrix between cancer cells (invasive tumor, proliferate invasive tumor, DCIS1 and DCIS2) and immune cells (CD4^+^ T cells, CD8^+^ T cells, IRF7^+^ DCs, LAMP3^+^ DCs and macrophages). To identify cell subtypes of DCIS, we used the attention vectors of DCIS1 (or DCIS2) on macrophages as cell-level features, and then calculated the PCA features. We used the top 50 PCs for K-means clustering and obtained two clusters as different subtypes. Differential genes for the two subtypes were then calculated based on the clustering results. For macrophages, we used the attention levels of DCIS1 (or DCIS2) as their cell-level features for hierarchical clustering and obtained two subtypes of macrophages. Differential genes were then computed based on the clustering results.

## Supplementary information

Supplementary Information is available for this paper.

## Acknowledgments

We thank all members of Eric and Wendy Schmidt Center as well as Cem Meydan, Ari Melnick and Eric Lander for their insightful comments on this work. We also thank Elvira Forte for scientific input and manuscript editing. Q.W. and L.Y. were supported by Eric and Wendy Schmidt Center Postdoctoral Fellowships. F.C. was supported by the Searle Scholars Award, the Burroughs Wellcome Fund CASI award, the Merkin Institute, and the NYSCF. F.C. is an NYSCF Roberston Investigator. R.X. was partially supported by NIDDK/NIH (5RC2DK135492-02). C.U. was partially supported by NCCIH/NIH (1DP2AT012345), NIDDK/NIH (5RC2DK135492-02), ONR (N00014-24-1-2687), DOE (DE-SC0023187), MIT J-Clinic for Machine Learning and Health, and the Eric and Wendy Schmidt Center at the Broad Institute.

## Declarations

### Data Availability

All datasets used for the training and evaluation of our model are publicly available. The HEST-1K [34] can be accessed at https://huggingface.co/datasets/MahmoodLab/hest. The Xenium human breast tissue slides were included in HEST-1K. We used the data version released by the original paper [37] (https://www.10xgenomics.com/products/xenium-insitu/preview-dataset-human-breast). The dataset of human lung tissues with pulmonary fibrosis [36] is deposited in the GEO database under accession number GSE250346.

### Code Availability

The code is available at https://github.com/uhlerlab/TissueFormer. The repository also contains a list of the required open-source packages with version numbers and instructions for reproducing the results and analyses performed in this study as well as a demo with instructions on how to apply the model to user-provided datasets.

### Author contribution

All authors designed the research. Q.W. developed the model and training pipeline. Q.W., Q.G., L.Y. and Z.L. performed data preprocessing and analysis. All authors wrote the paper.

## Footnotes

1 For technologies with subcellular resolution (e.g. Xenium), each note in our model corresponds to a cell, while for spot-level technologies (e.g. Visium), each node is a spot and thus we use a spot-centered image patch and spot-level gene expression.

2 ST refers to a spatial transcriptomics technology which has lower resolution than Visium.

3 We used the cell type annotations provided by the dataset [36].

