## Supplementary material for "Multi-Modal Foundation Model with Whole-Slide Attention Enables Transferrable Digital Pathology at Single-Cell Resolution": figs. S1-20

### Supplementary Figures

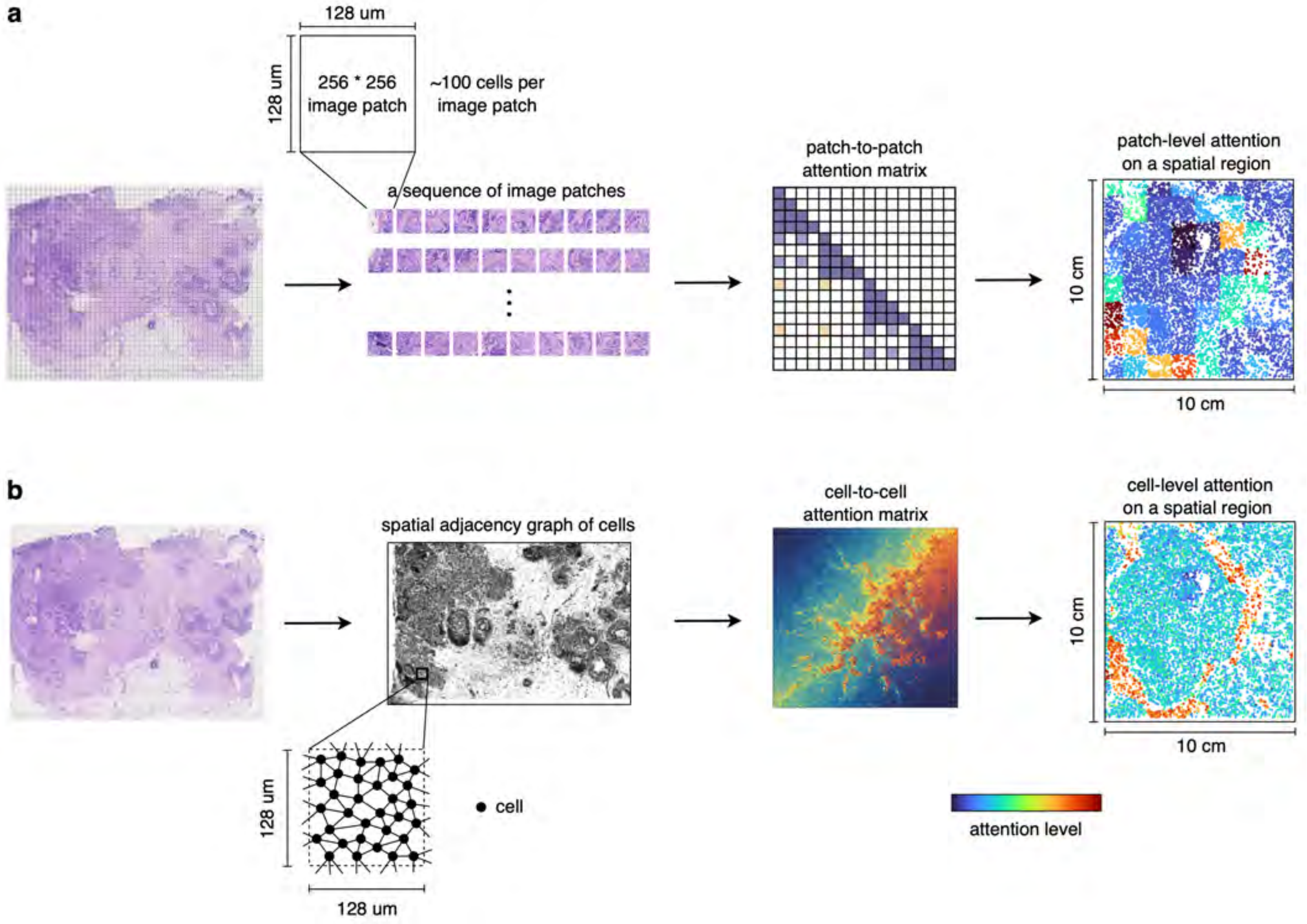


**fig. S1: Comparison of patch-to-patch attention (Prov-GigaPath [24]) and cell-to-cell attention (TissueFormer). a,** Illustration of Prov-GigaPath's (and other state-of-the-art models') patch-level representation and attention: the model first partitions a whole-slide image into a sequence of nonoverlapping image patches, each containing roughly a hundred cells, and then computes the attention between image patches. The resulting patch-level attention only captures coarse-grained interaction since the cellular features are averaged across all cells within an image patch. **b,** Illustration of TissueFormer's cell-level representation and attention: the model represents each single cell through a spatial adjacency graph and computes the cell-to-cell attention across the whole slide. Our cell-level attention framework offers a scalable approach for fine-grained interrogation of cellular dependencies.

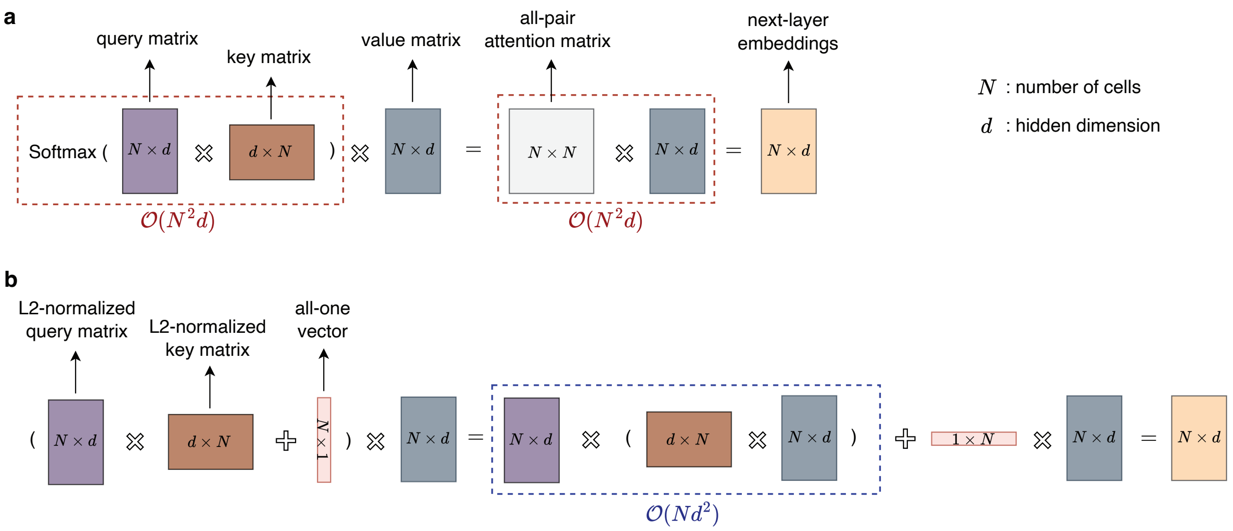


**fig. S2: Illustration for the Softmax attention model and the scalable attention model. a,** Computation flow for softmax-based attention that requires quadratic computational (i.e., $\mathcal{O}(N^2d)$) complexity with respect to the number of cells. **b,** Computation flow for the proposed scalable attention model that reduces computational complexity to the linear order (i.e., $\mathcal{O}(Nd^2)$) with respect to numbers of cells ($N$), without sacrificing the expressivity for capturing all-pair influence.

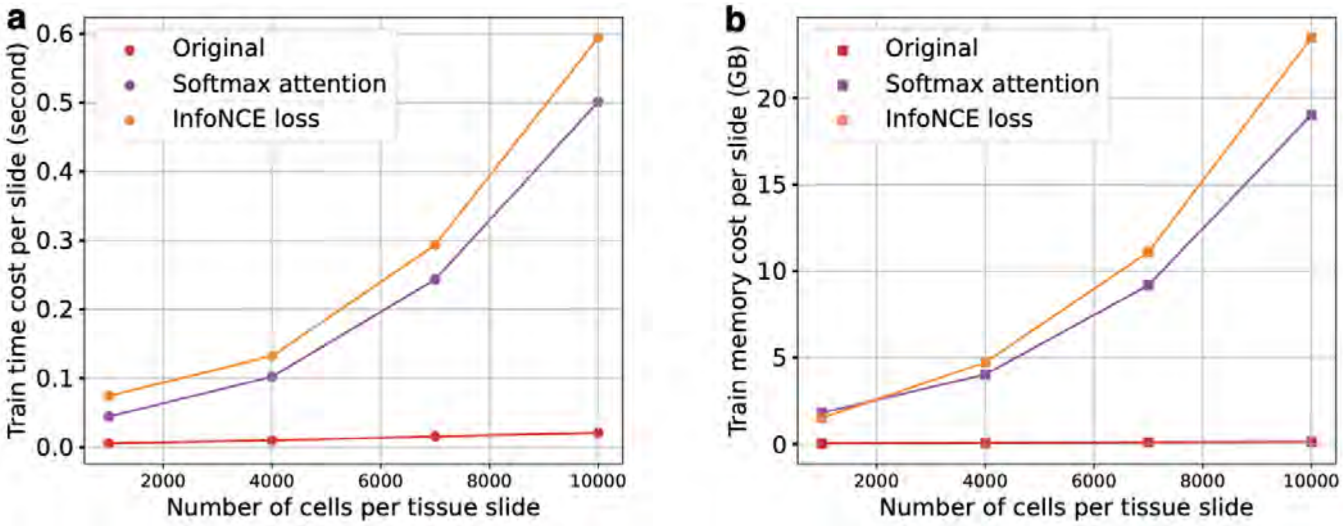


**fig. S3: Comparison of time and memory costs. a,** Plots showing the average training time per epoch per slide of TissueFormer, the model variant replacing our proposed attention with standard Softmax attention [38], and the model variant replacing our scalable contrastive loss with standard InfoNCE loss [55]. **b,** Plots showing the maximum GPU memory cost per slide for these models during training.

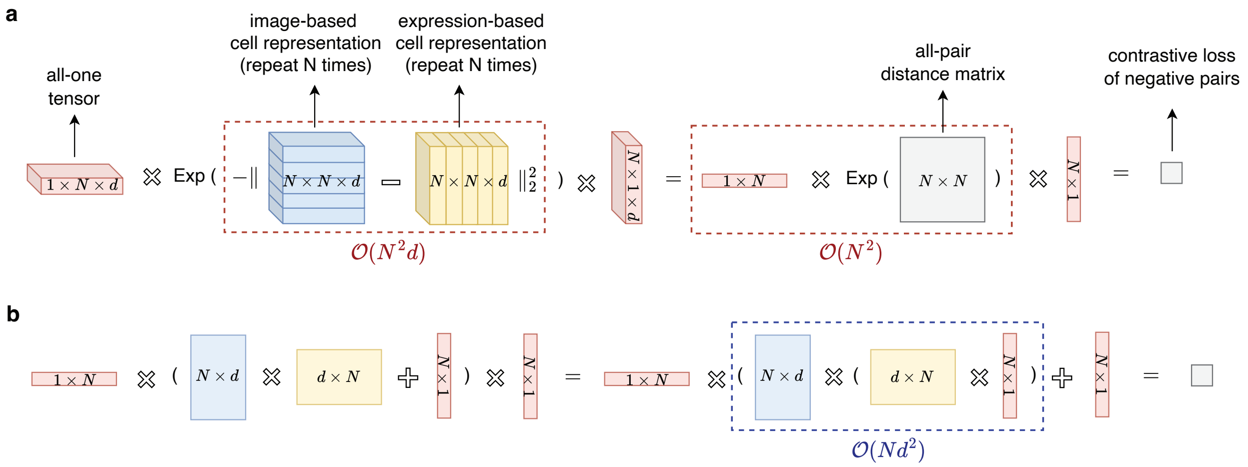


**fig. S4: Illustration for the standard contrastive loss and the scalable contrastive loss. a,** Computation flow for InfoNCE-based contrastive loss where the all-pair distance matrix requires quadratic complexity (i.e., $\mathcal{O}(N^2d)$). **b,** Computation flow for the proposed scalable contrastive loss that allows efficient computation within linear order (i.e., $\mathcal{O}(Nd^2)$) with respect to numbers of cells ($N$).

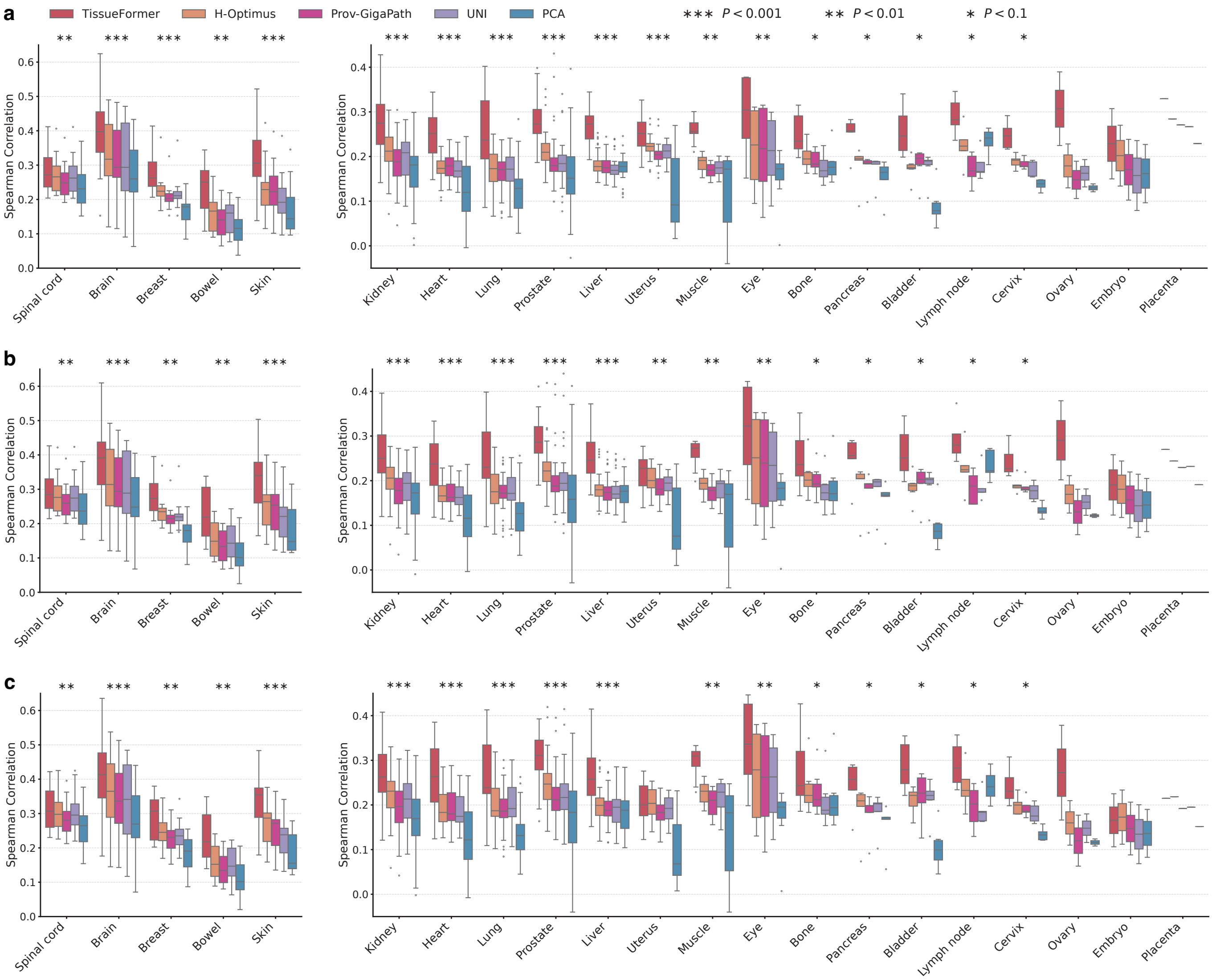


**fig. S5: Comparison of gene expression prediction under the protocol of generalization across organs. a-c,** Box plots showing the Spearman correlation between the predicted expression and ground-truth expression of top 4000 (**a**), 2000 (**b**) and 1000 (**c**) highly variable genes, respectively. The Spearman correlation is first computed per spot across evaluated genes and then averaged across spots of one slide. Each box shows the distribution of Spearman correlation scores of test tissue slides from each organ. The P-value indicates the significance level that TissueFormer outperforms the best competitor in each case, with Wilcoxon signed-rank test.

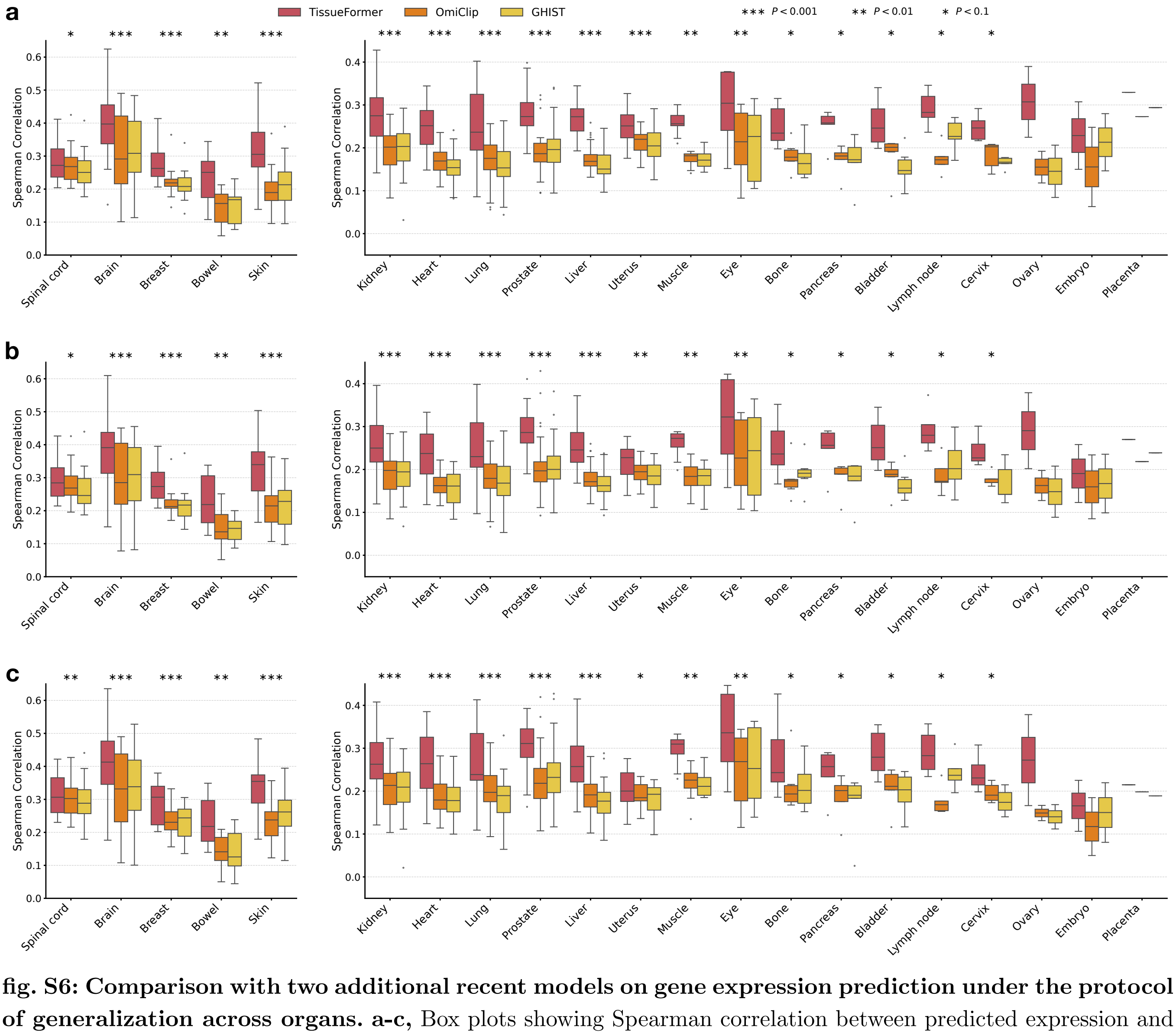


**fig. S6: Comparison with two additional recent models on gene expression prediction under the protocol of generalization across organs. a-c,** Box plots showing Spearman correlation between predicted expression and ground-truth expression of the top 4000 (**a**), 2000 (**b**) and 1000 (**c**) highly variable genes, respectively.

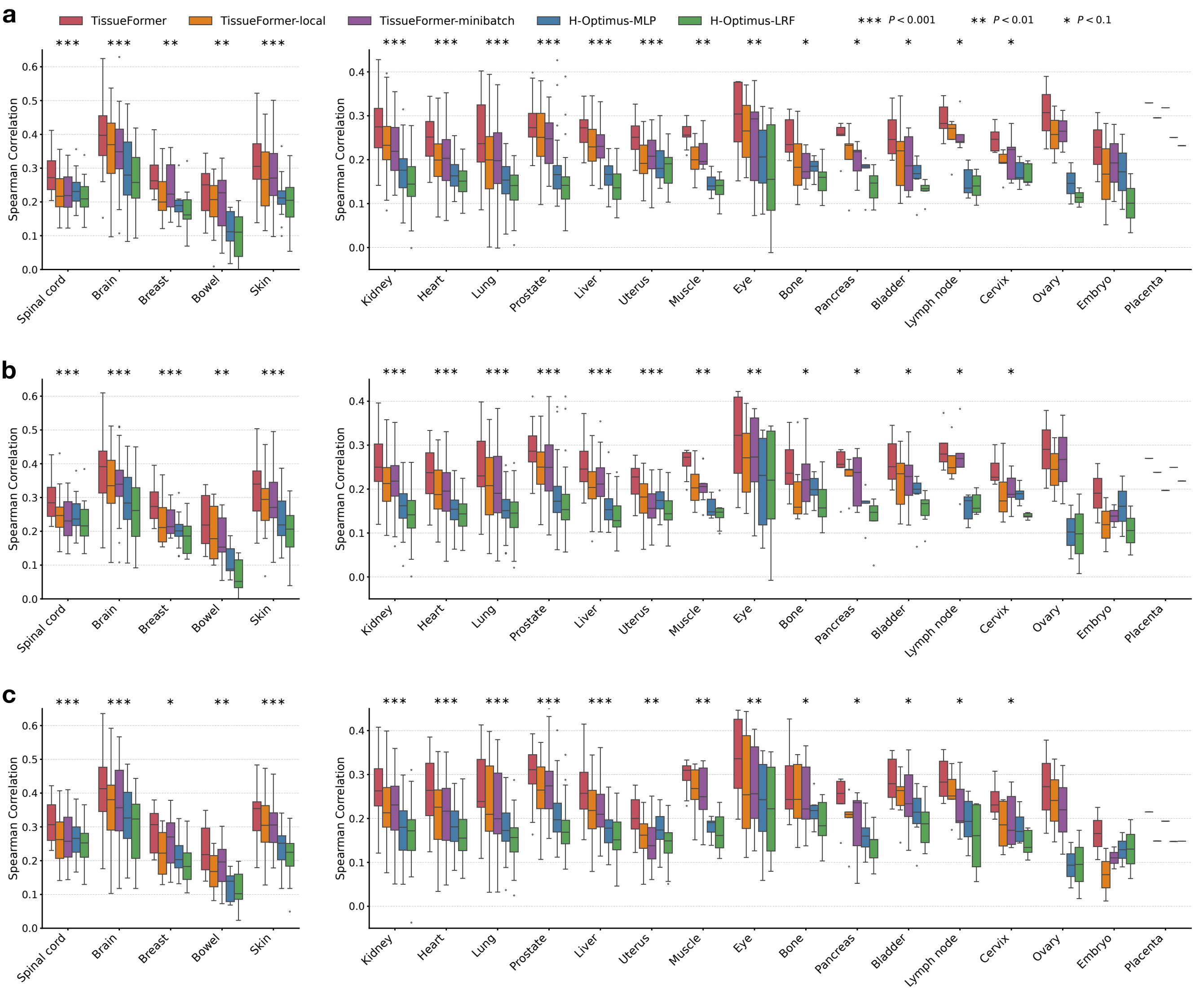


**fig. S7: Comparison with multiple model variants on gene expression prediction under the protocol of generalization across organs.** TissueFormer-local replaces the global attention of TissueFormer with the local counterpart that only attends to neighboring cells, and TissueFormer-minibatch adopts mini-batch training that takes a subset of cells as input. H-Optimus-MLP replaces the linear regression predictor with a shallow MLP, and H-Optimus-LRF takes the embeddings of five nearest neighbor patches as input for MLP for prediction. **a-c**, Box plots showing Spearman correlation between predicted expression and ground-truth expression of the top 4000 (**a**), 2000 (**b**) and 1000 (**c**) highly variable genes, respectively.

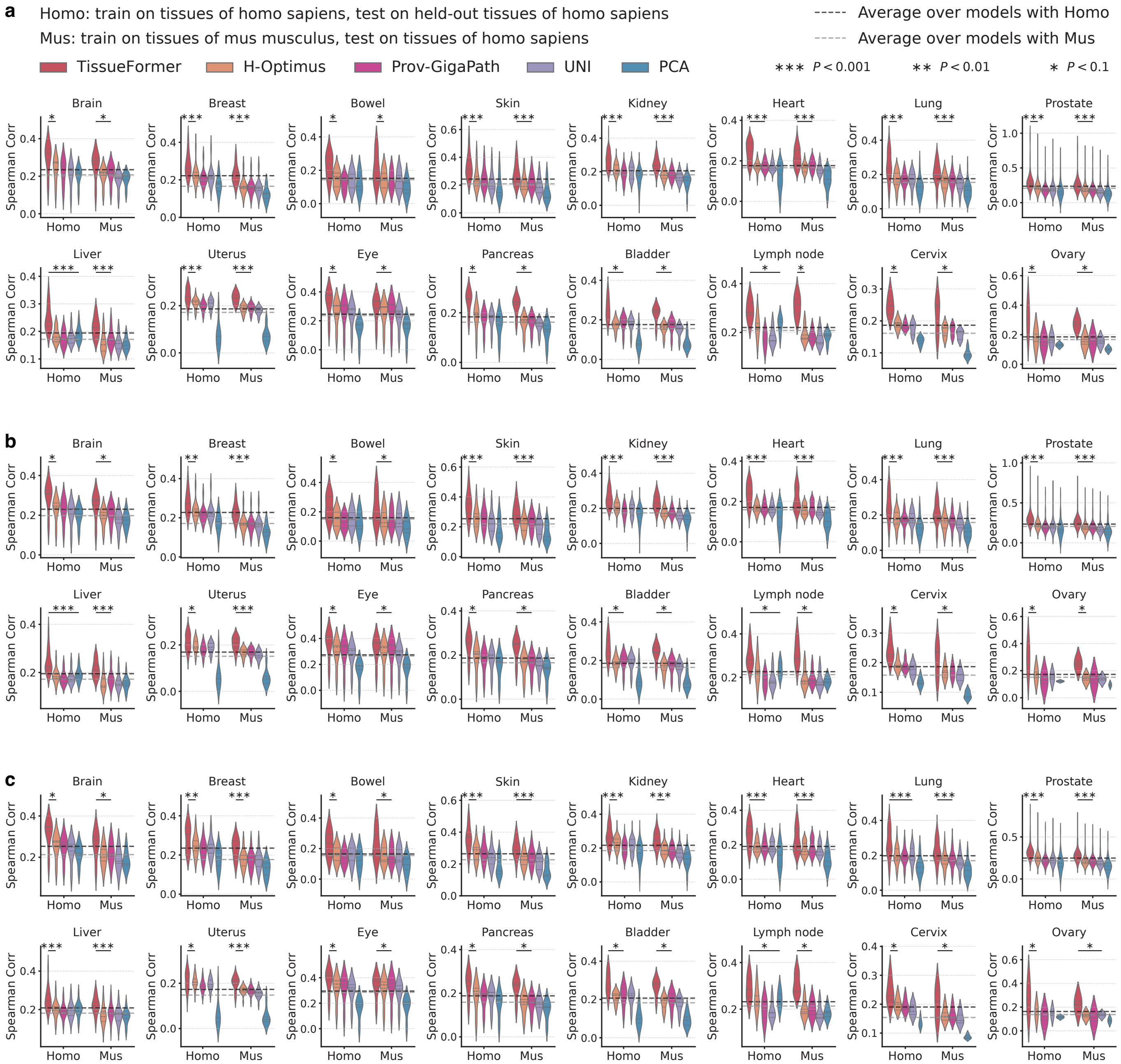


**fig. S8: Comparison of gene expression prediction under the protocol of generalization across species. a-c,** Violin plots showing the Spearman correlation between the predicted expression and ground-truth expression of top 4000 (**a**), 2000 (**b**) and 1000 (**c**) highly variable genes, respectively. The Spearman correlation is first computed per spot across evaluated genes and then averaged across spots of one slide. Each violin shows the distribution of Spearman correlation scores of test tissue slides from one organ, and each plot compares the model performance in two settings, *Homo sapiens* (Homo) and *Mus musculus* (Mus). The horizontal dashed lines show the averaged performance of all models. The P-value indicates the significance level that TissueFormer outperforms the best competitor in each case, with Wilcoxon signed-rank test.

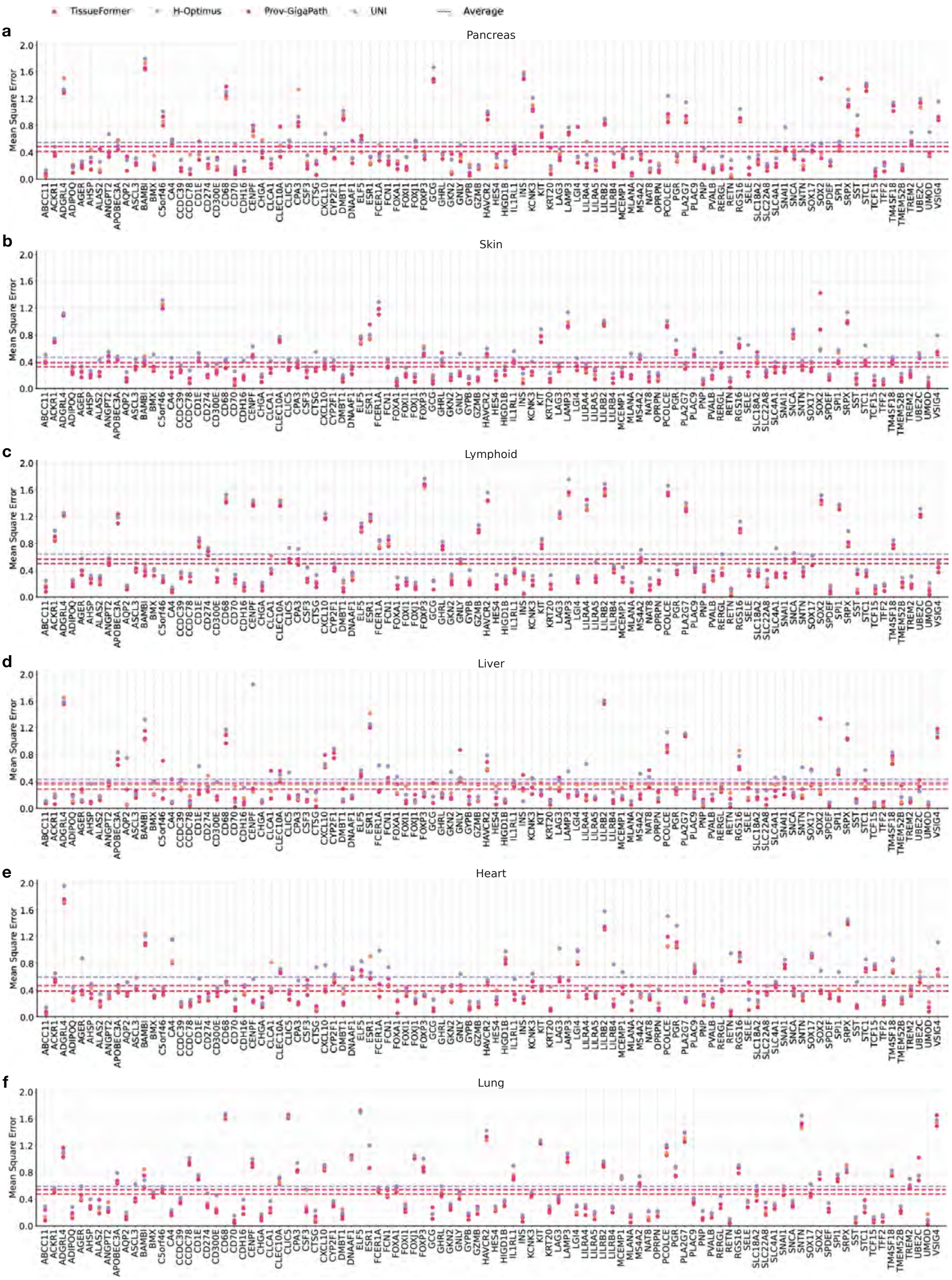


**fig. S9: Gene-wise evaluation of gene expression prediction at single-cell resolution. a-f,** Plots showing gene-wise mean square error (MSE) between ground-truth expression and the predicted expression by TissueFormer and competing methods across top 100 highly-variable genes of test Xenium slides from six organs. The dotted horizontal lines show the averaged MSE of different models across all genes.

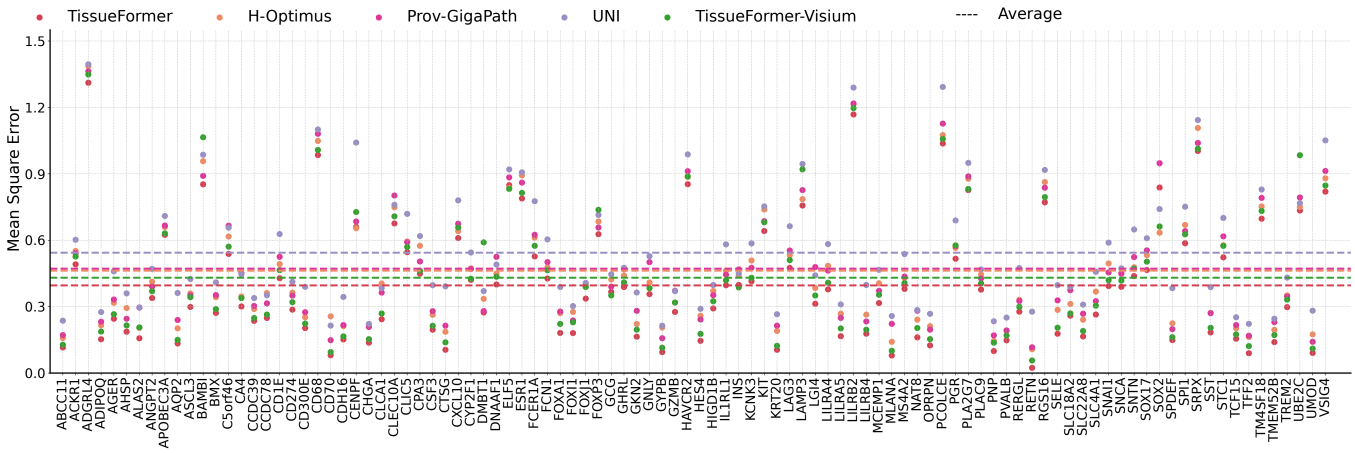


**fig. S10: Comparison with TissueFormer trained only on the Visium slides (TissueFormer-Visium) for gene expression prediction on Xenium slides.** Plots showing gene-wise Mean Squared Error (MSE) between ground-truth expression and predicted expression by different methods across the top 100 highly-variable genes of Xenium slides from six organs. The dotted horizontal lines show the averaged MSE of different models across all genes.

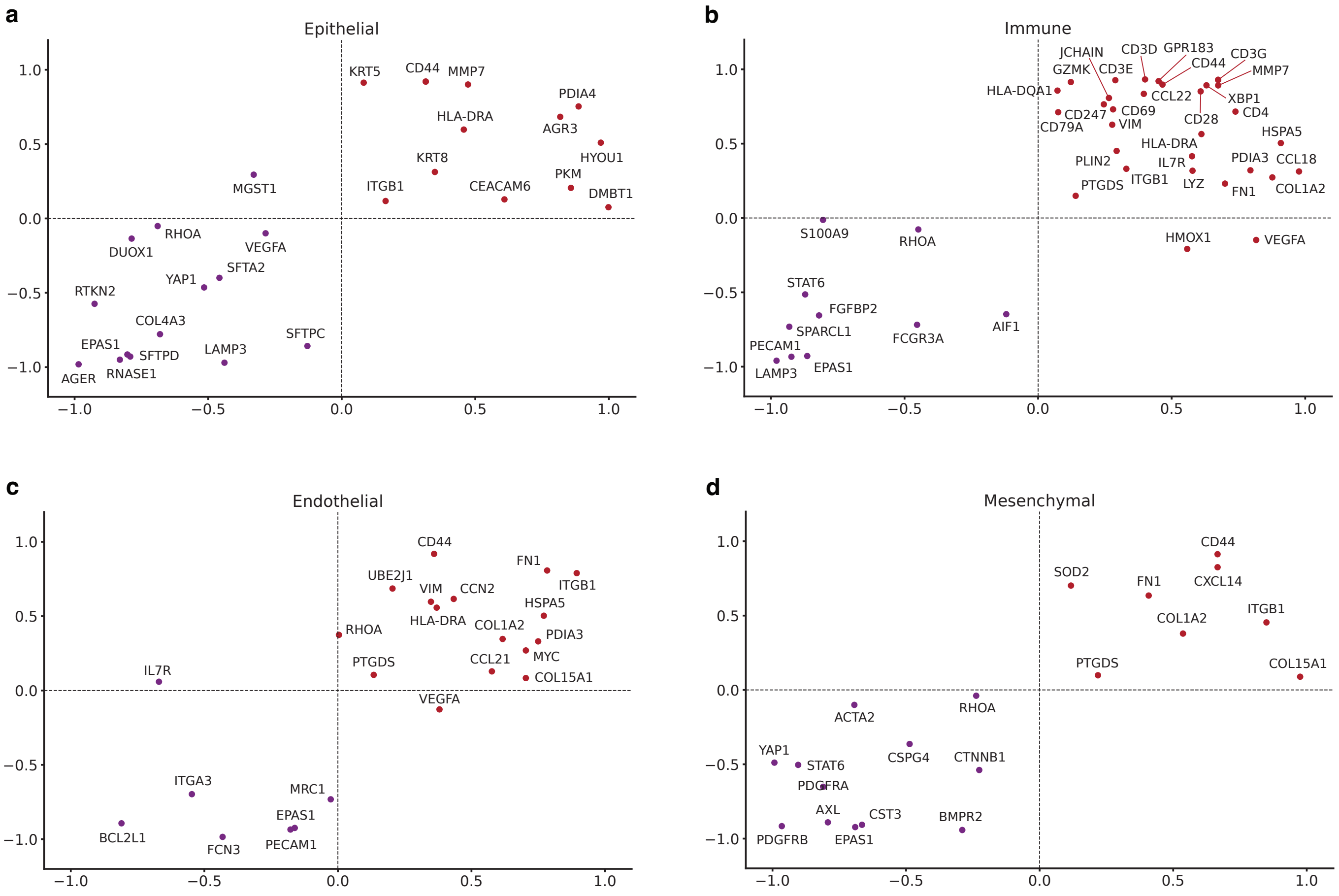


**fig. S11: Evaluation of TissueFormer's gene expression prediction for disease-associated markers. a-d,** Plots showing the consistency between the prediction and the ground-truth expression of gene markers that exhibit significantly ($P < 0.01$) positive or negative correlation with pathology scores of lung tissues with pulmonary fibrosis at cell-type level. In each plot, the Pearson correlation between ground-truth expression and pathology scores (x-axis) is compared with the Pearson correlation between predicted expression and pathology scores (y-axis).

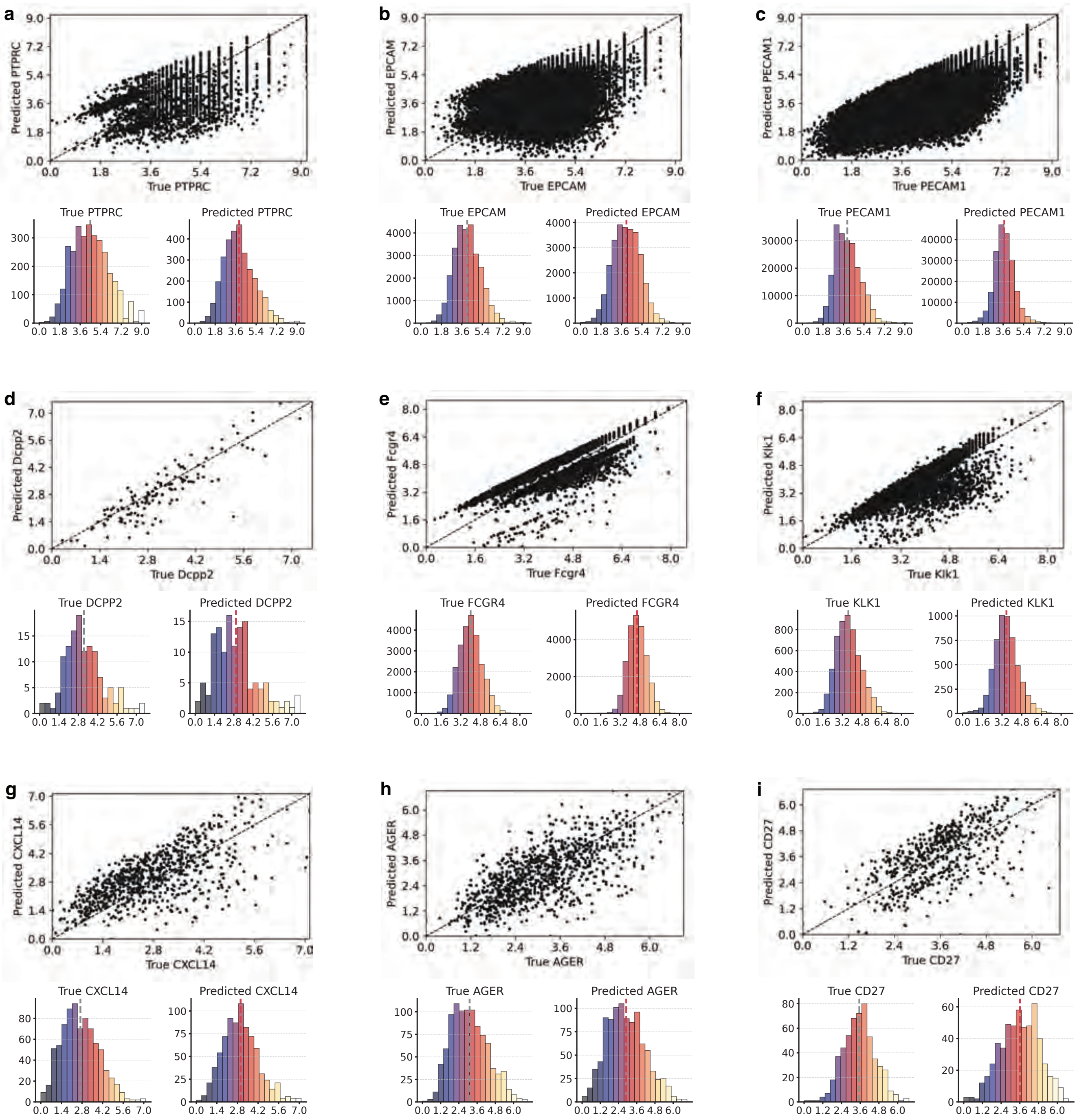


**fig. S12: Demonstration of TissueFormer's prediction of marker genes' expression. a-i,** Scatter plots showing the predicted and true expression levels for marker genes on tissue slides from the test set, and histogram plots comparing the distributions of the predicted and true expression. On the first row (a-c): marker genes for common cell types; second row (d-f): marker genes from bone tissues that are unseen in training; third row (g-i): marker genes strongly associated with pulmonary fibrosis.

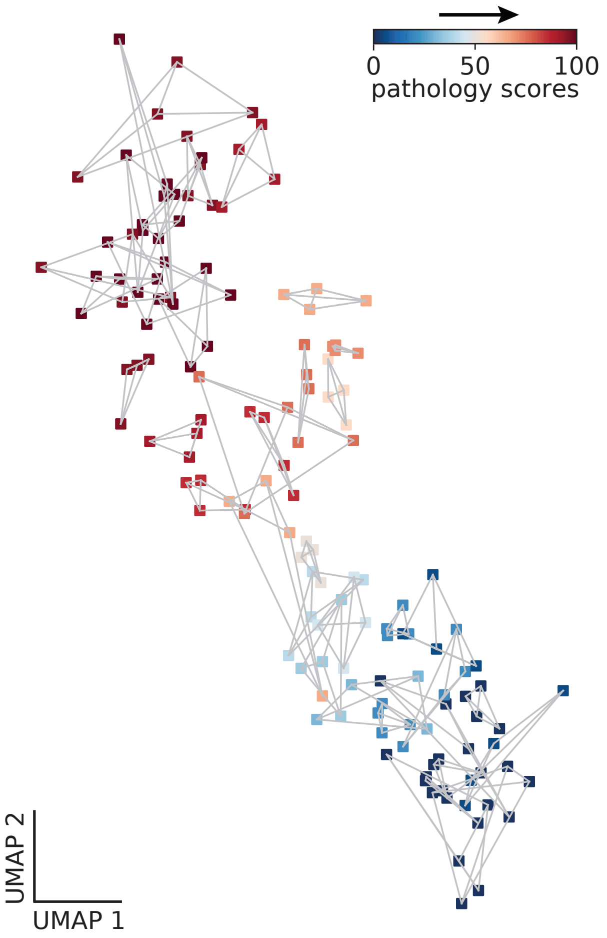


**fig. S13: UMAP embeddings of cropped sections from pulmonary fibrosis tissue slides**. We split each slide into four cropped sections (top left, top right, bottom left and bottom right) and compute the embeddings of sections by aggregating cell-level representations of cells within each section. The cropped sections from the same slide are connected by lines and share the same slide-level pathology score.

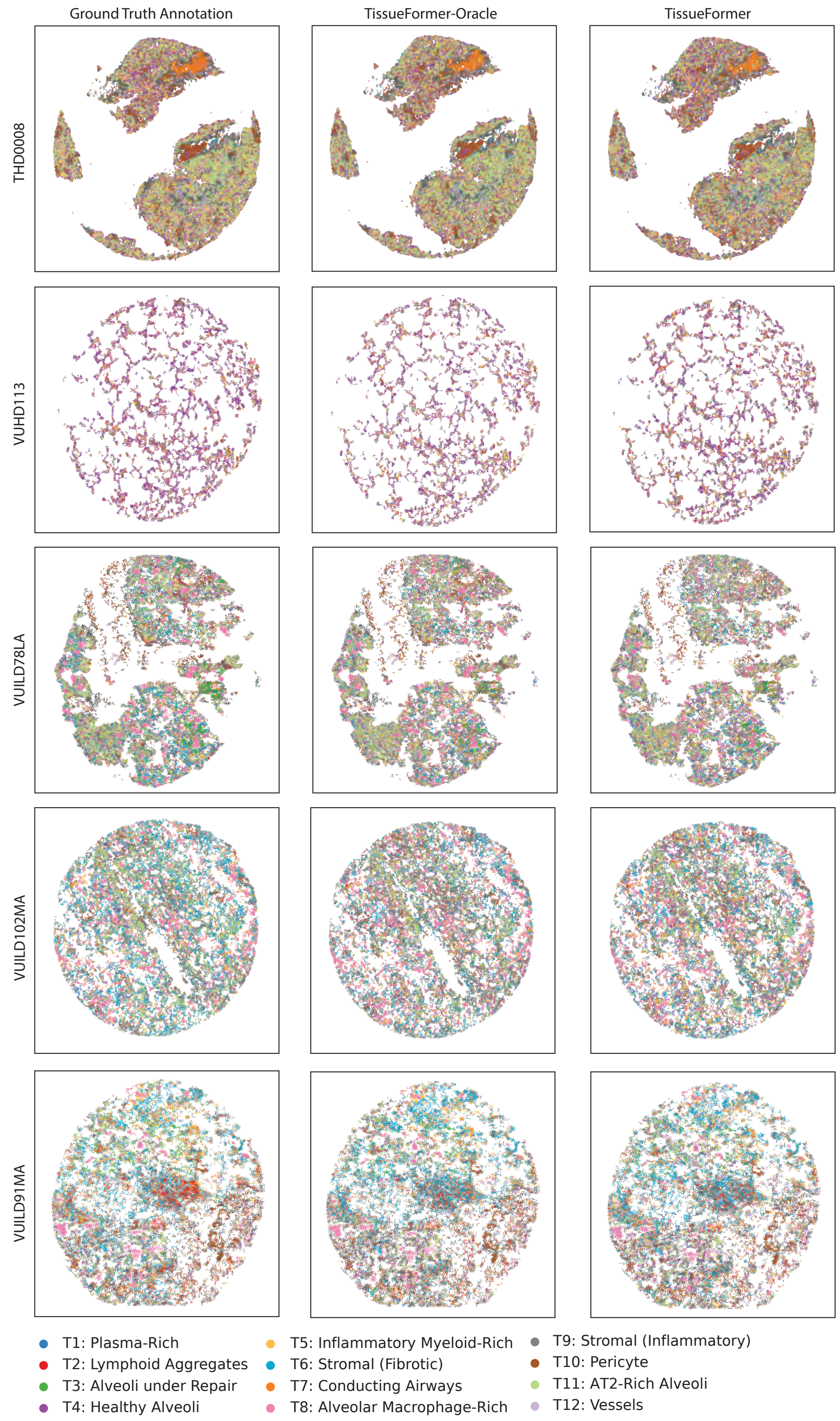


**fig. S14: Spatial niches classification on tissue slides from pulmonary fibrosis samples.** Images showing the ground-truth annotation of spatial niches (computed by transcriptomics profiles and neighboring information of cells) and the predicton by TissueFormer-Oracle and TissueFormer. Tissue slides include unaffected (THD0008 and VUHD113), less affected (VUILD78LA) and more affected (VUILD102MA and VUILD91MA) samples from [36].

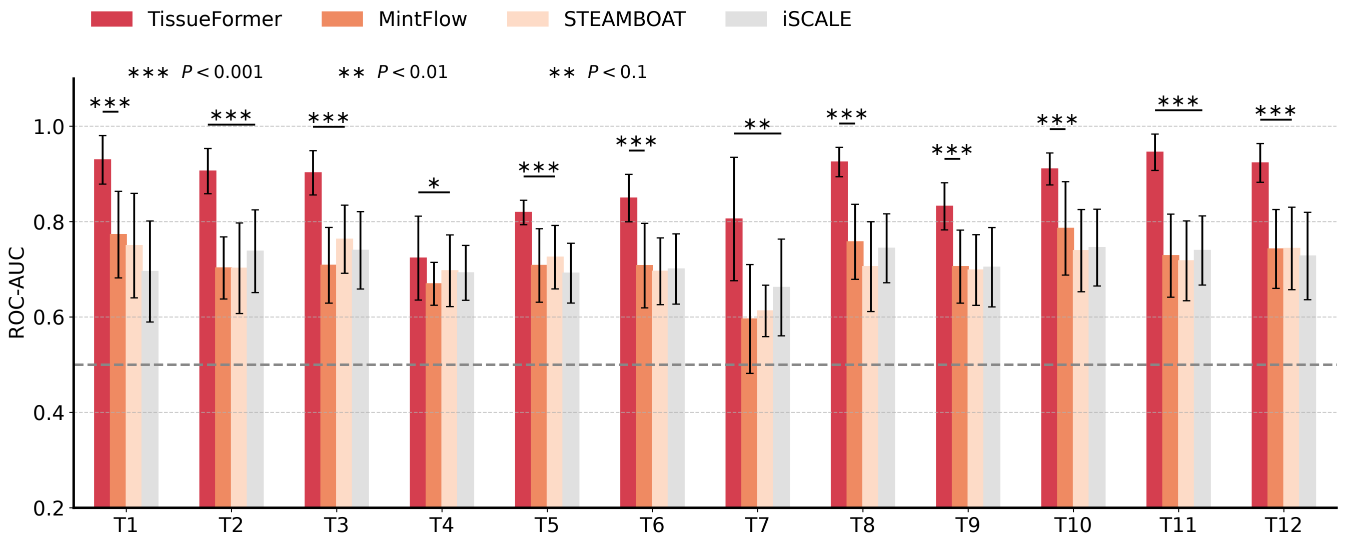


**fig. S15: Comparison with three additional recent models on spatial niche classification.** Bar plots showing the ROC-AUC scores under 12 spatial niche labels. The horizontal dotted line marks ROC-AUC of 0.5 that corresponds to random guessing. The P-values indicate the significance level at which TissueFormer outperforms the best competitor in each case, based on Wilcoxon signed-rank test.

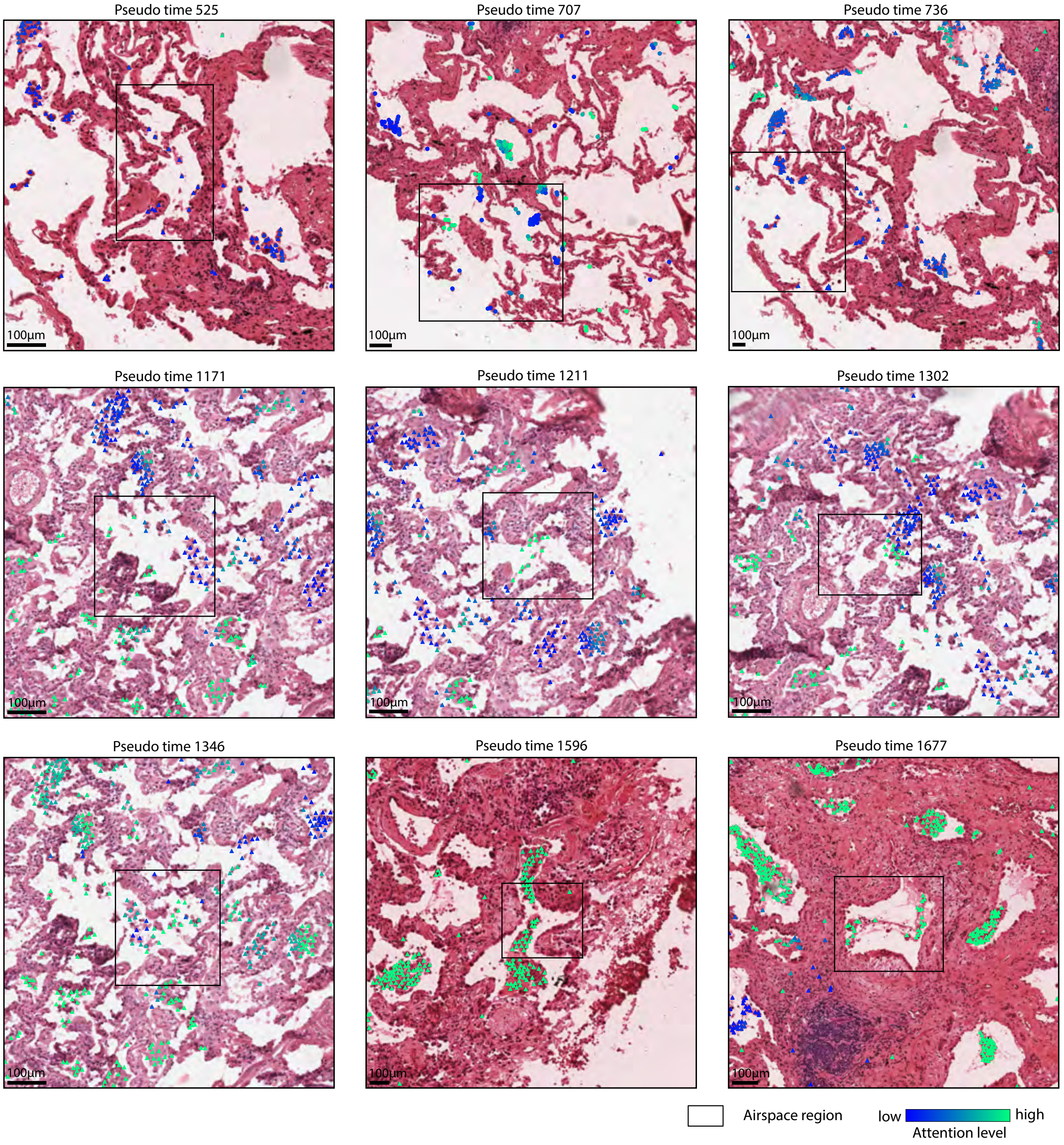


**fig. S16: Representative examples of attention maps on airspace regions of pulmonary fibrosis tissue slides from different pseudo times.** Overlaid to the H&E images are the alveolar macrophages. The colors indicate the attention levels that each alveolar macrophage receives from activated fibrotic fibroblasts.

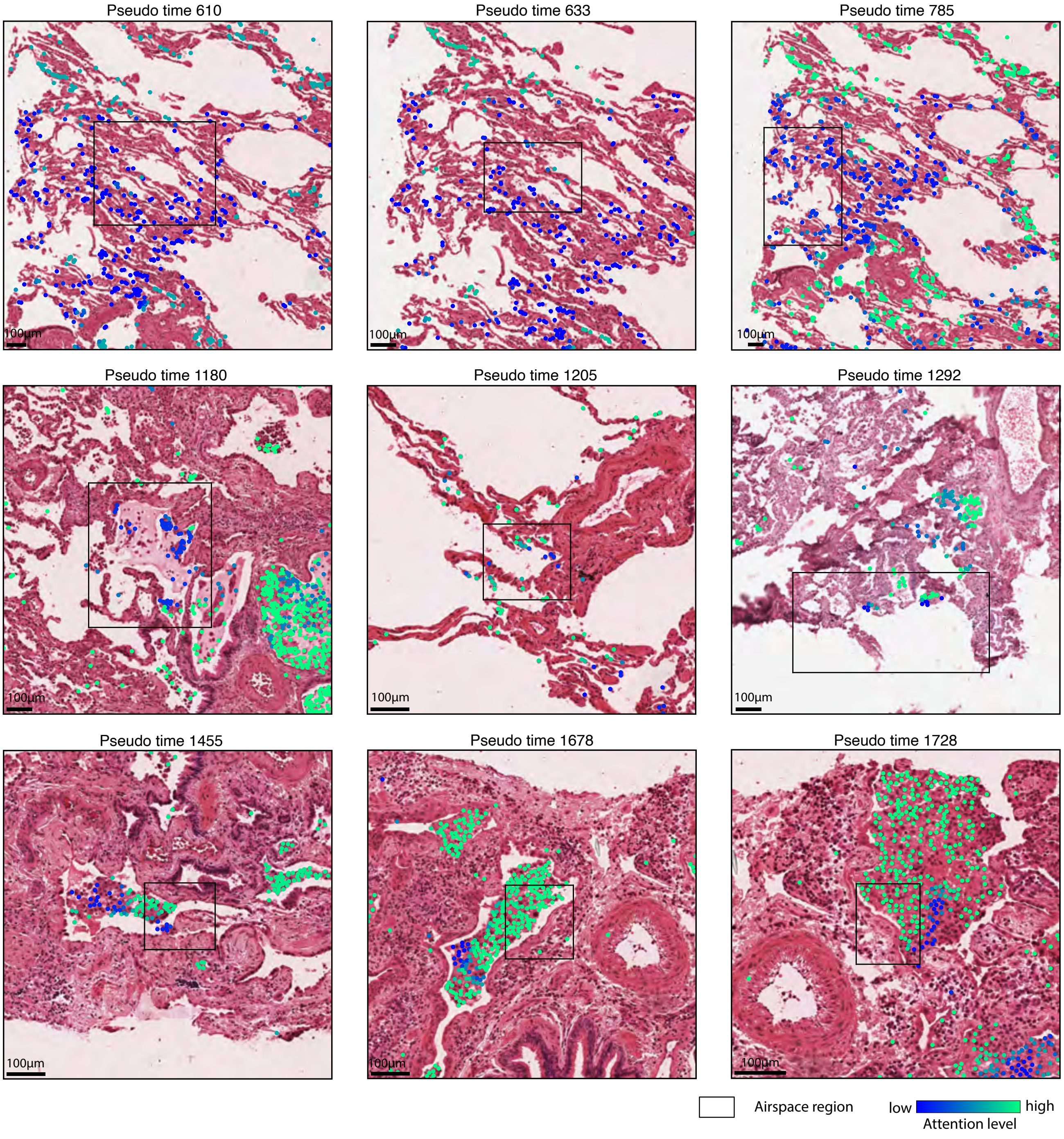


**fig. S17: Representative attention maps on airspace regions in pulmonary fibrosis tissue slides from different pseudo times.** Overlaid on the H&E images we show the SPP1$^+$ macrophages. The colors indicate the attention levels that each SPP1$^+$ macrophage receives from activated fibrotic fibroblasts.

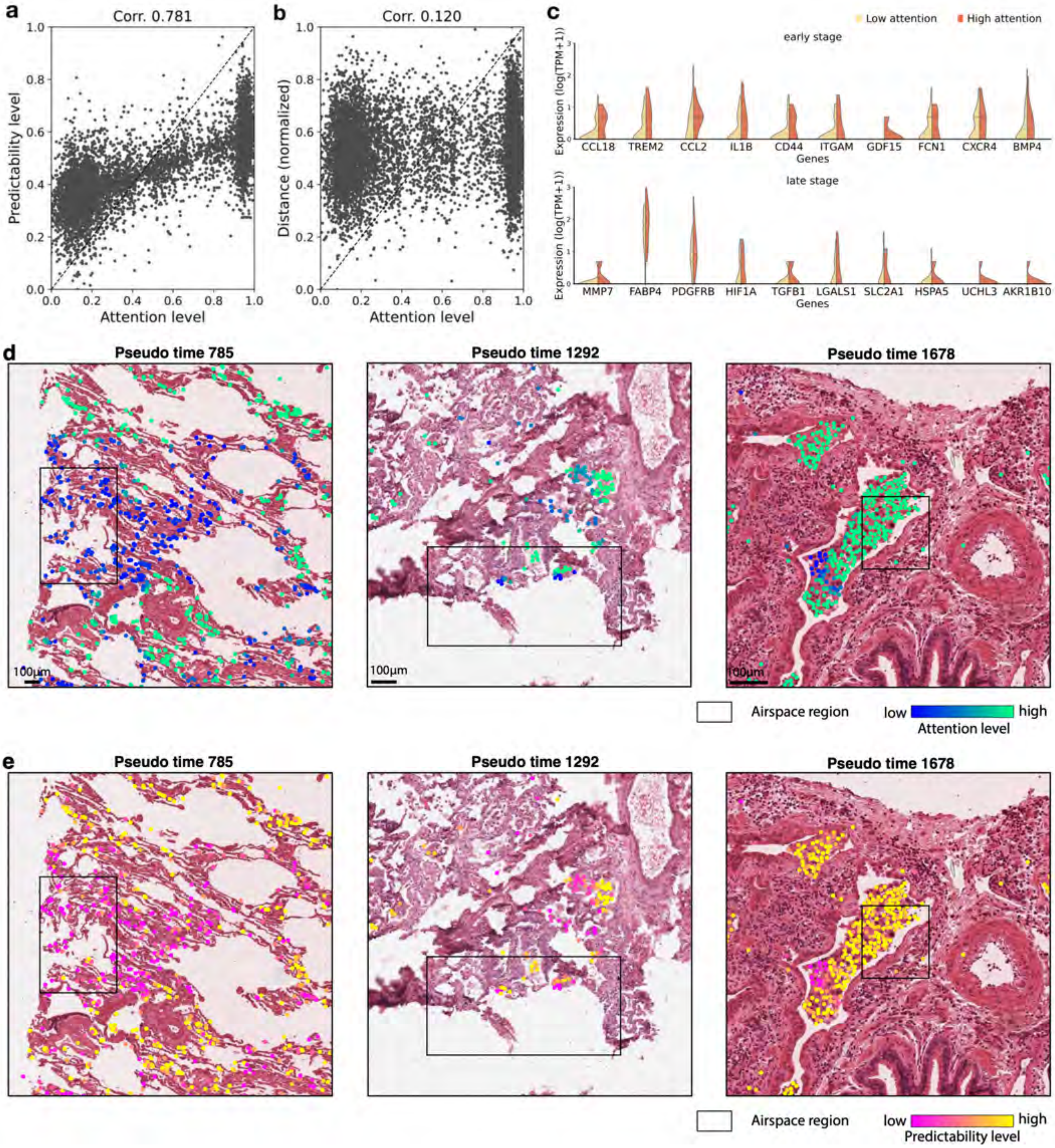


**fig. S18: Interpretation of cellular attention maps. a,** Plot showing the correlation between attention levels and predictability levels of SPP1$^+$ macrophages. For any given SPP1$^+$ macrophage, we compute its predictability level by: 1) using this macrophage's image embedding for predicting gene expression of activated fibrotic fibroblasts in the airspace region of interest, 2) computing the Spearman correlation between predicted and measured expression across all genes, 3) averaging the Spearman correlations of activated fibrotic fibroblasts in the airspace of interest. **b,** Plot showing the Spearman correlation between attention levels and normalized distance (from activated fibrotic fibroblasts in the airspace of interest) of SPP1$^+$ macrophages. **c,** Top ten differentially expressed genes of SPP1$^+$ macrophages with high and low attention levels from activated fibrotic fibroblasts. The attention levels above (resp. below) the 50th percentile are treated as high (resp. low) attention. **d-e,** Representative examples of the correlation between attention levels and predictability levels of SPP1$^+$ macrophages. **d,** Overlaid on the H&E images we show the SPP1$^+$ macrophages (defined based on paired transcriptomic data). The colors indicate the attention levels that each SPP1$^+$ macrophage receives from activated fibrotic fibroblasts. **e,** Overlaid on the same H&E images, SPP1$^+$ macrophages are colored by the predictability levels with regard to activated fibrotic fibroblasts.

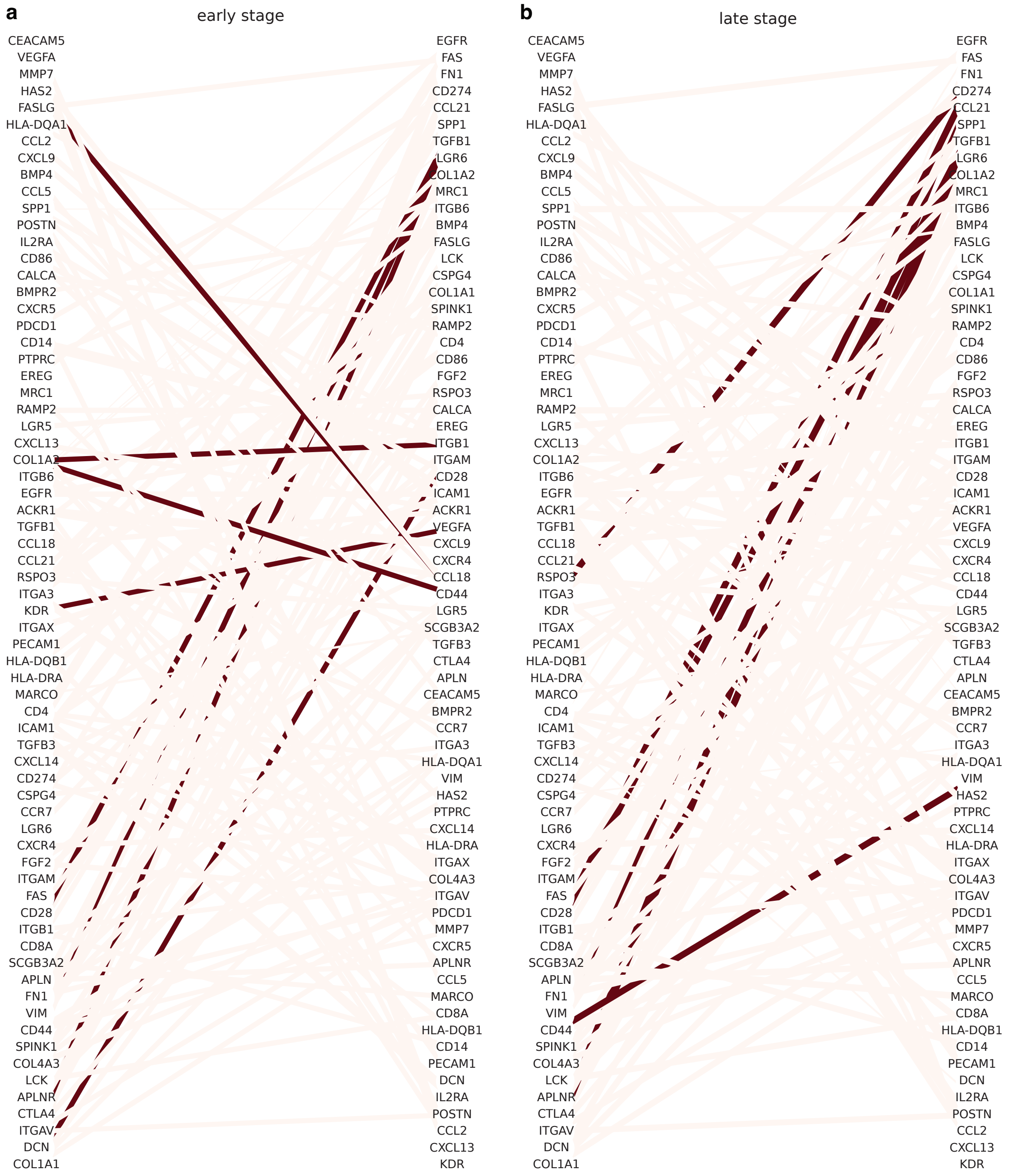


**fig. S19: Ligand-receptor pairs between SPP1$^+$ macrophages and activated fibrotic fibroblasts.** In each bipartite graph, the left and right columns represent genes measured in macrophages and fibroblasts, respectively. Dark-colored edges mark the LR pairs whose co-expression levels are significantly associated with attention levels (FDR $> 0.01$). Light-colored edges represent non-significant LR pairs.

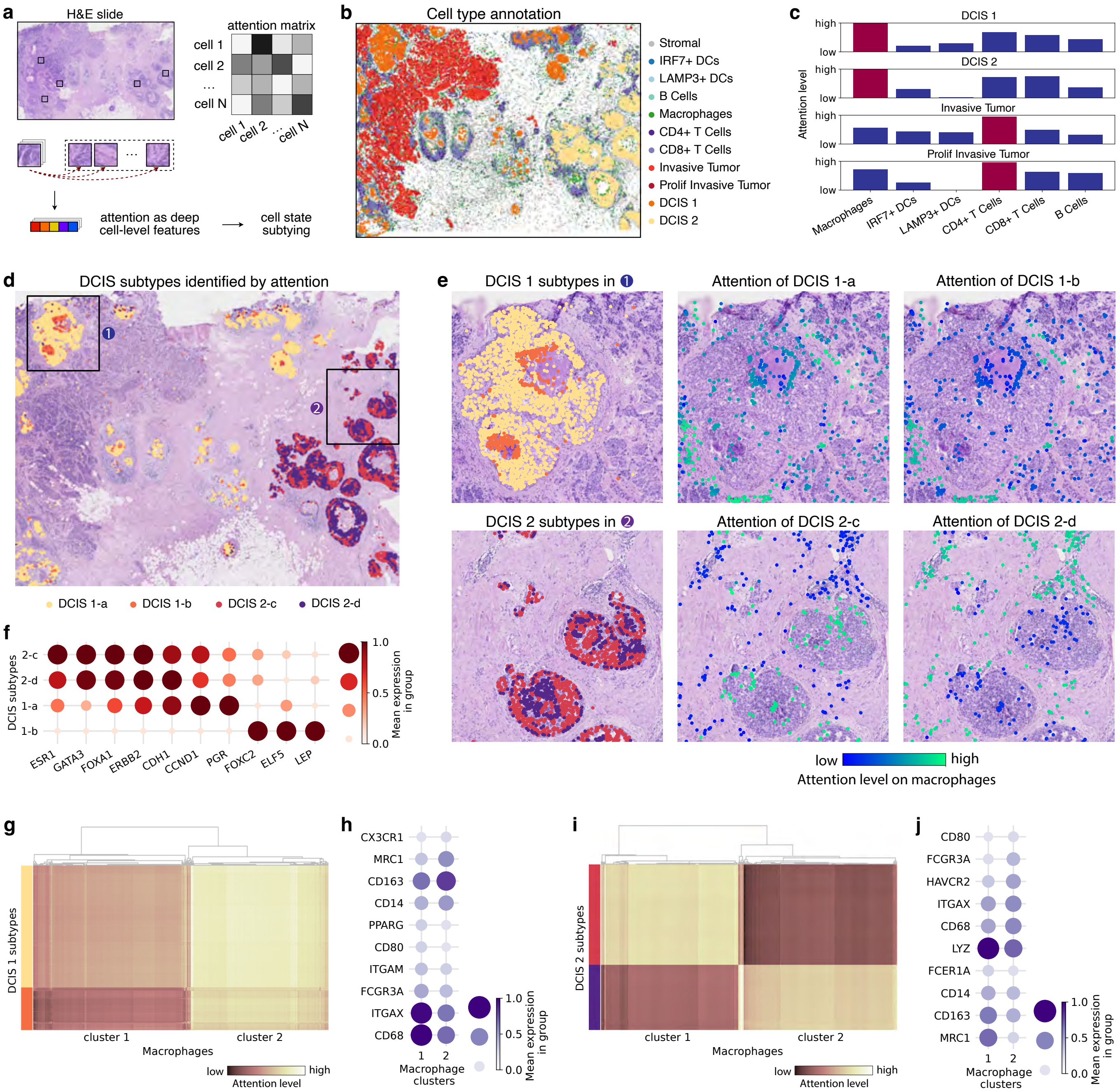


**fig. S20: Attention-based modeling of cell-level features for cell subtyping.** **a,** Illustration of the analysis on breast tumor tissue slides, where we extracted the pairwise attention from the pretrained TissueFormer on H&E images as cell-level features. The latter is further used for subtyping cells with different states. **b,** Example of cell type annotation of a tissue slide. **c,** Averaged attention levels of tumor cells on different immune cells, per tumor stage. **d,** H&E image overlaid with colors indicating the spatial distributions of four DCIS subtypes (two from DCIS 1 and two from DCIS 2) defined by unsupervised clustering based on attention levels of DCIS 1 (and DCIS 2, respectively) on macrophages. **e,** Images showing the magnified region (1) (top row) and region (2) (bottom row) defined in **d**. The leftmost images indicate the DCIS 1 and DCIS 2 subtypes. The central and right images indicate the geometric mean of attention levels of a DCIS subtype on macrophages within the region. **f,** Top differential expressed genes between the two subtypes of DCIS 1 and two subtypes of DCIS 2. We ranked DCIS subtypes from the most malignant (top) to the most benign ones (bottom) according to the expression levels of these differential genes. **g,** Heatmap showing the attention levels of DCIS 1 on macrophages, which are grouped into two clusters according to different attention patterns of two DCIS 1 subtypes. **h,** Top differential genes of two macrophage clusters in **h**. **i,** Heatmap showing the attention levels of DCIS 2 on macrophages, which are grouped into two clusters according to different attention patterns of two DCIS 2 subtypes. **j,** Top differential genes of the two macrophage clusters in **i**.

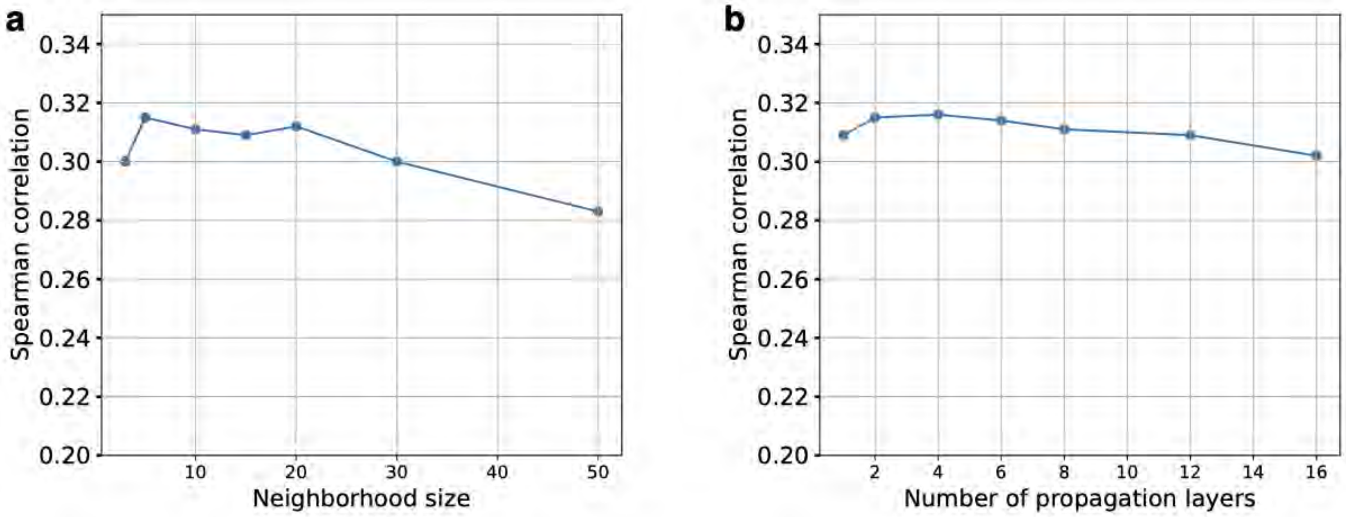


**fig. S21: Performance impact of hyper-parameters including neighborhood size (a) and number of propagation layers (b).** Plots showing the average Spearman correlation for gene expression prediction under the protocol of generalization across organs.
